# Interpretable spatial multi-omics data integration and dimension reduction with SpaMV

**DOI:** 10.1101/2025.08.02.668264

**Authors:** Yang Liu, Kexin Ma, Haoran Xu, Ke Xu, Yunfei Hu, Zhenhan Lin, Jiangli Lin, Bo Han, Shuaicheng Li, Zhixiang Lin, Xin Maizie Zhou, Lu Zhang

## Abstract

Spatial multi-omics technologies have revolutionized our understanding of biological systems by providing spatially resolved molecular profiles from multiple perspectives. Existing spatial multi-omics integration methods often assume that data from different modalities share a common underlying distribution, aiming to project them into a single unified latent space. This assumption, however, can obscure the unique insights offered by each modality, thereby limiting the full potential of multi-omics analyses. To address this limitation, we present the Spatial Multi-View (SpaMV) representation learning algorithm, which captures both the shared information across modalities and the distinct, modality-specific information, enabling a more comprehensive and interpretable representation of spatial multi-omics data. Through extensive evaluation on both simulated and real-world datasets, SpaMV demonstrates superior spatial domain clustering performance and provides users with more interpretable dimension reduction for downstream analysis. Moreover, SpaMV effectively annotates cell types within clusters of a mouse thymus dataset, highlighting its effectiveness in interpretable dimensionality reduction.

---

Spatial multi-omics technologies revolutionize the study of biological systems by integrating genomics, transcriptomics, proteomics, and metabolomics within their native tissue architecture [1, 2]. By capturing multi-omics information, these technologies enable a comprehensive view of cellular states, which have the potential to enhance the accuracy of spatial segmentation in complex tissues [3]. For instance, combining spatial transcriptomics and proteomics reveals drug resistance mechanisms (e.g., SPP1+ macrophage-CAF interactions) and metastatic drivers at tumor boundaries or tertiary lymphoid structures. These insights uncover dynamic communication networks and immunosuppressive hubs, providing actionable targets to disrupt tumor progression and therapy resistance [4].

Recent advancements in spatial multi-omics technologies have driven the development of integration algorithms to tackle the complexities of combining diverse omics datasets. SpatialGlue [5] employs graph convolutional networks (GCNs) to extract modality-specific representations, which are integrated via an attention mechanism to form a unified latent space for unsupervised clustering. CellCharter [6] uses neural networks to derive spot-level representations, aggregating them with neighboring spots to capture local spatial contexts and enhance integration. COSMOS [7] also leverages GCNs for feature extraction, incorporating a deep graph infomax model and spatial regularization to ensure spatially coherent embeddings. MISO [8], treating H&E images as an additional modality, extracts features using neural networks and applies element-wise matrix multiplication between modality embeddings to model cross-modal interactions before concatenation.

Existing spatial multi-omics integration algorithms typically project multi-modal data into a shared latent space, which can result in the loss of modality-specific information when the data across modalities do not fully overlap. For example, chromatin accessibility peaks may be primed for future transcriptional activation without corresponding gene expression, as demonstrated in [9]. Such modality-specific signals, unique to certain omics layers, are critical for understanding complex biological processes. Losing these signals can compromise the accuracy of downstream analyses, such as cellular state characterization. Therefore, an effective integration framework must preserve both shared cross-modal information and modality-specific features to fully capture the richness of spatial multi-omics data.

Here, we present Spatial Multi-View representation learning (SpaMV), a novel spatial multi-omics integration algorithm designed to explicitly disentangle cross-modal shared features and modality-specific private features into distinct latent spaces. Building on variational autoencoders (VAEs) [10], SpaMV incorporates three key components to achieve robust decomposability: (1) a dual-encoder architecture, where each omics modality employs separate encoders, i.e, graph attention networks (GATs) to capture spatially informed shared and private signals, coupled with a modality-specific decoder for faithful reconstruction; (2) auxiliary measurement models, which are utilized to minimize mutual information between inferred private latent variable from one modality and the data from other modalities, thereby preventing leakage of shared information into the private latent spaces; and (3) non-parametric independence test to enforce statistical independence between shared and private latent variables, avoiding contamination of shared space by modality-specific signals. SpaMV provides two types of output, i.e., spatial domain clustering and interpretable dimension reduction, to enable comprehensive analysis and understanding of the spatial multi-omics data.

We evaluated SpaMV against state-of-the-art spatial multi-omics integration methods, including SpatialGlue [5], CellCharter [6], COSMOS [7], SMOPCA [11], MISO [8], and spaMultiVAE [12], using three simulated and four real-world spatial multi-omics datasets spanning a broad range of omics combinations, including epigenome-transcriptome, H3K27ac-H3K27me3 histone modifications, transcriptome-metabolome, and transcriptome-proteome integrations. Quantitative evaluations on both synthetic and manually annotated data demonstrated that SpaMV consistently outperformed existing methods. In addition, we present the interpretable biological topics and their modality-specific feature associations, highlighting SpaMV’s ability to reveal coherent and biologically meaningful patterns across multiple modalities. Notably, by leveraging these modality-specific topics, SpaMV accurately annotated cell types in mouse thymus tissue—a task that could not be reliably achieved by current integration methods.

## 1 Results

### The overview of SpaMV

SpaMV adopts a multi-view assumption for the multi-modality data (Fig. 1a), positing that each omics modality ***x***_*i*_ can be viewed as a different perspective of the same tissue slice, encompassing both cross-modal shared information ***z***_*s*_ and modality-specific private information ***z***_*pi*_ [13–15]. The structure of SpaMV is shown in Fig. 1b. In the SpaMV model, each omics dataset is processed using two encoders to extract both shared and private latent representations, along with a decoder to reconstruct the original values. SpaMV utilizes GAT as encoders to consider the spatial correlation between spots. The underlying graph for the GAT is constructed by leveraging both spatial proximity and data similarity: an edge is established between two nodes if they are either adjacent in physical space or exhibit high similarity in their observation. By incorporating both spatial and data-based similarities, the GAT enables SpaMV to adaptively capture both local and global correlations among spots. SpaMV concatenates the shared and private representations derived from the same omics data ***x***_*i*_, using them as the input to the decoder to reconstruct ***x***_*i*_. This process is referred to as self-modality reconstruction. To ensure that shared latent variables effectively capture necessary shared information, we designed a cross-modality reconstruction process, which reconstructs omics data ***x***_*i*_ using shared representations inferred from another omics data ***x***_*j*_. Besides, to prevent the leakage of shared information into the inferred private latent variables, we introduced additional multi-layer perceptrons named measurement models to minimize the mutual information between inferred private latent variables ***z***_*pi*_ and other omics data ***x***_*j*_. Lastly, to prevent the leakage of private information into the inferred shared latent variables, SpaMV minimizes the dependency between inferred shared and private latent variables using a non-parametric kernel-based independence test, i.e., Hilbert-Schmidt independence criterion (HSIC) [16]. SpaMV integrates the shared representations from different omics by a mixture of experts operator [17], and outputs all private and shared representations for down-stream tasks such as spatial domain clustering and interpretable dimension reduction (Fig. 1c). Full derivation and technical details of SpaMV are provided in Methods and Spplementary Note. B.

**Fig. 1:**
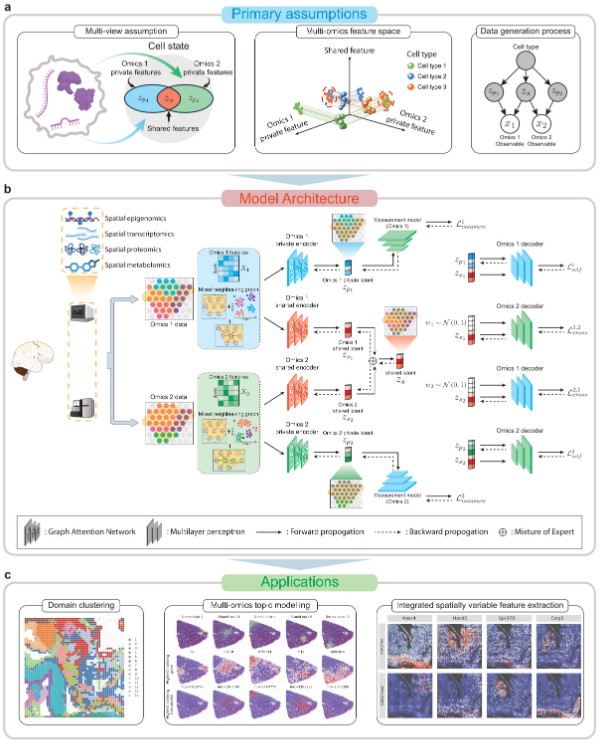
The assumptions, architecture, and applications of SpaMV. **a**, SpaMV operates under the assumption that each omics dataset, denoted as ***X***_*i*_, is generated from a combination of unobserved shared latent variables ***z***_*s*_ and private latent variables ***z***_*pi*_. These latent variables are influenced by the unobserved cell state. Relying solely on partial information can adversely5affect the performance of downstream tasks such as domain clustering. **b**, SpaMV employs two distinct GAT encoders for each omics dataset to extract its shared and private information, respectively. The shared representations are efficiently integrated using a mixture of experts operation. SpaMV implements self-modality reconstruction to enhance the capture of private information in the inferred private representations, and cross-modality reconstruction to bolster shared information in the inferred shared representations. Additionally, SpaMV utilizes measurement models to prevent shared information from infiltrating the inferred private representations and incorporates a non-parametric independence test (HSIC) to inhibit private information from contaminating the inferred shared representations. **c**, SpaMV uses inferred latent variables to produce downstream analysis, including domain clustering, interpretable multi-omics topic modeling, and integrated spatial variable feature extraction.

SpaMV provides two operational modes tailored to different tasks. For domain clustering, SpaMV utilizes multi-layer perceptrons as the decoder for each omics dataset, concatenating the shared and private representations to serve as inputs for the down-stream clustering algorithm. For interpretable dimension reduction, SpaMV adopts a matrix multiplication decoder, which reconstructs the omics data ***x***_*i*_ by the product of the latent variables [***z***_*s*_, ***z***_*pi*_] and a feature embedding matrix *β*_*i*_. In this framework, each dimension of [***z***_*s*_, ***z***_*pi*_] corresponds to an interpretable biological topic, while each column of *β*_*i*_ quantifies the contribution of each feature in ***x***_*i*_ to a particular topic. Higher values indicate greater feature importance for the respective topic. We also refer to this task as topic modeling throughout the paper.

### Benchmarking SpaMV and competing algorithms on simulated data

To validate the effectiveness of SpaMV, we compared its performance against other spatial multi-omics integration algorithms using three simulated datasets characterized by diverse patterns (Methods). In these datasets, we defined several shared and private factors based on their presence across different omics data. For example, there is a private factor specific to omics 1 and another specific to omics 2, indicating that these factors are exclusive to omics 1 or omics 2, respectively (Fig. 2a). In contrast, the other seven factors are present in both omics datasets.

**Fig. 2:**
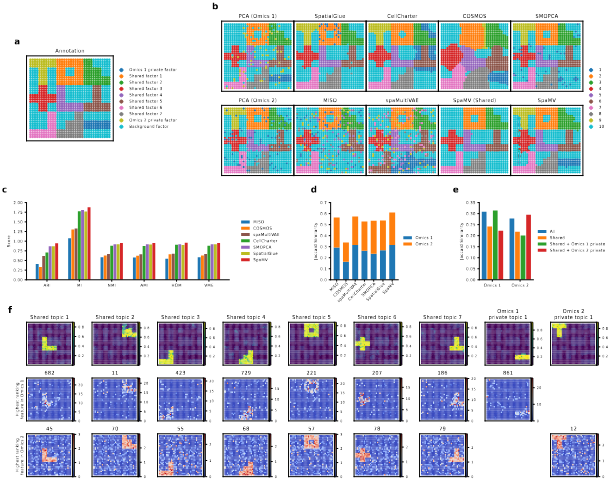
SpaMV accurately decomposes shared and private features in simulation. **a**, Ground truth of the simulated data 1. **b**, Spatial plots of clustering results: first column shows the mclust clustering on PCA embeddings from omics 1 and omics 2; subsequent columns display clustering results from SpatialGlue, CellCharter, COSMOS, SMOPCA, MISO, spaMultiVAE, and SpaMV. **c**, Quantitative evaluation of the evaluated algorithms using six supervised metrics. **d**, Jaccard similarity scores for competing algorithms and SpaMV, where the Jaccard similarity scores for SpaMV were computed using the learned shared and omics-specific private embeddings. **e**, Jaccard similarity scores for SpaMV using different embeddings. **f**, Spatial plot of the learned topics and the highest ranking features identified by SpaMV. ARI: Adjusted Rand Index, MI: Mutual Information, AMI: Adjusted Mutual Information, NMI: Normalized Mutual Information, HOM: Homogeneity, VME: V-measure.

We began by assessing the domain clustering performance of SpaMV alongside eight other single-omics and multi-omics integration algorithms. As shown in Fig. 2b, SpaMV was the only algorithm capable of identifying all factors. Other algorithms, such as SpatialGlue, CellCharter, and SMOPCA, successfully identified all factors present in the Omcis 2 dataset but failed to recover the omics 1 private factor, implying a bias towards the omics 2 modality and a neglect of signals in omics 1.

The supervised scores for spatial domain clustering results (Fig. 2c) demonstrated that SpaMV achieved superior clustering performance across all metrics. Additionally, to investigate whether the inferred private latent variables captured modality-specific information, we employed the Jaccard similarity as an unsupervised metric. Notably, concatenating SpaMV’s omics 1 private representation with the shared representation resulted in a higher Jaccard similarity for omics 1, suggesting that the private embedding contains complementary information specific to omics 1 that is not present in the shared embedding (Fig. 2e). In contrast, concatenating the omics 2 private representation with the shared embedding decreased the Jaccard similarity for omics 1, indicating that the omics 2 private embedding does not contribute additional information about omics 1. These results confirm the effective decomposability of the SpaMV model. Furthermore, SpaMV achieved better Jaccard similarity scores relative to other algorithms for both omics datasets (Fig. 2d).

Finally, we applied SpaMV’s interpretable topic modeling module to visualize the learned topics and their associated highest ranking features (Fig. 2f). The spatial distributions of both shared and private topics closely aligned with the corresponding ground truth spatial factors, demonstrating SpaMV’s ability to disentangle biologically meaningful structures. Importantly, the top-ranked features associated with each topic accurately reflected the corresponding spatial patterns, further validating the interpretability of the learned representations. To assess the robustness and generalizability of SpaMV, we repeated the same analysis on two additional simulated datasets with different spatial patterns (Extended Data Figs. 1 and 2). The consistent alignment between predicted topics, feature relevance, and ground truth across all datasets confirms SpaMV’s robustness and its ability to generalize across varying spatial multi-omics contexts.

### SpaMV enhances spatial multi-omics integration in mouse embryo spatial epigenome-transcriptome data

We applied SpaMV on a real-world dataset from an embryonic day 13 (E13) mouse embryo, which includes co-profiling of the spatial transcriptome and epigenome [18]. For anatomical reference, we utilized the EMAGE mouse embryo spatial gene expression database [19] to annotate major anatomical regions, including the thalamus, lateral ventricle, and spine (Fig. 3a). The visualized results (Fig. 3c) reveal that SpaMV successfully identifies all key anatomical structures in the E13 mouse embryo, including thalamus (cluster 3), hindbrain (cluster 6), spine (cluster 7), wall of diencephalon (cluster 10), trigeminal ganglion (cluster 12), and choroid plexus (cluster 13). In contrast, other methods demonstrated notable limitations: SpatialGlue failed to distinguish the wall of diencephalon, CellCharter conflated the lateral ventricle with the thalamus, SMOPCA merged the lateral ventricle and wall of diencephalon, while COSMOS and MISO erroneously split the hindbrain into two different clusters. To quantitatively benchmark performance, we evaluated all six algorithms using the domain annotations from the original study (Fig. 3b). Across all metrics, SpaMV consistently outperformed other algorithms. Additionally, the Jaccard similarity scores (Fig. 3e-f) confirmed that SpaMV more effectively preserves modality-specific information, reinforcing its strength in integrative representation learning.

**Fig. 3:**
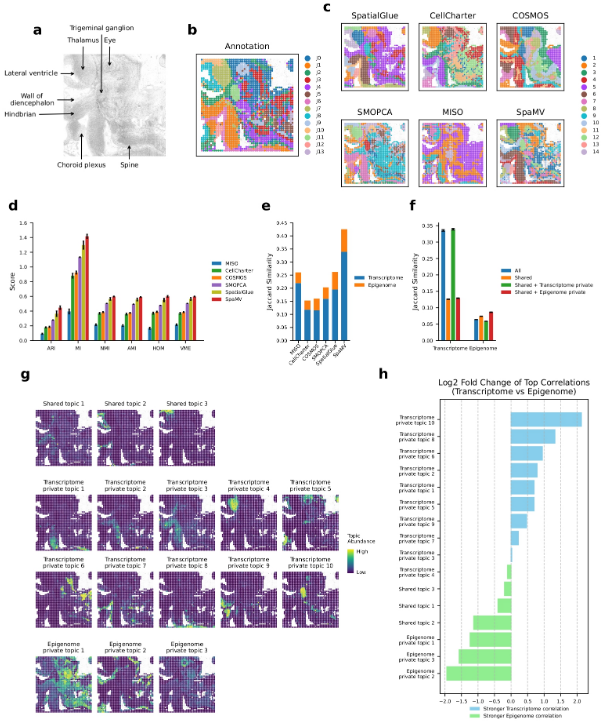
Interpretable SpaMV analysis of spatial transcriptome-epigenome mouse embryo data. **a**, Image with manual annotation of the mouse embryo data. **b**, Domain annotations as described in [18]. **c**, Domain clustering results from six spatial multi-omics integration algorithms: SpatialGlue, CellCharter, COSMOS, SMOPCA, MISO, and SpaMV. **d**, Mean and standard deviation of performance for the six evaluated algorithms, based on six supervised metrics over 10 repetitions. **e**, Jaccard similarity scores for competing algorithms and SpaMV, where the Jaccard similarity scores for SpaMV were computed using the learned shared and omics-specific private embeddings. **f**, Jaccard similarity scores for SpaMV using different embeddings **g**, Spatial plots illustrating the topics learned b8y SpaMV using the topic modeling mode. **h**, Log2 fold change of the mean Pearson correlation coefficients between transcriptome and epigenome for each topic, calculated from the top 10 correlated genes. ARI: Adjusted Rand Index, MI: Mutual Information, AMI: Adjusted Mutual Information, NMI: Normalized Mutual Information, HOM: Homogeneity, VME: V-measure.

To further explore the shared and private information within this spatial multiomics data, we employed the interpretable topic modeling mode of SpaMV. In this analysis, chromatin accessibility data were transformed into gene activity scores (GASs) (Methods). Our SpaMV method enabled the identification of 16 distinct topics (Fig. 3g), including 3 shared topics, 10 transcriptome private topics, and 3 epigenome private topics. The prevalence of transcriptome-specific topics indicates that chromatin accessibility is less effective in distinguishing domains within this dataset compared to transcriptomics, consistent with previous findings reported by [18].

Specifically, the shared topics predominantly encompass regions such as the lateral ventricle (shared topic 2) and the central nervous system (shared topics 1 and 3). Transcriptome private topics mainly include the hindbrain (transcriptome private topics 1, 2, and 3), thalamus (transcriptome private topic 4), ventral part of medulla oblongata (transcriptome private topics 7 and 8), and the trigeminal ganglion (transcriptome private topic 10). In contrast, epigenome private topics primarily cover the wall of the diencephalon (epigenome private topic 2) and eye regions (epigenome private topic 3). Furthermore, the topic-specific top-ranking genes identified through the learned feature embeddings showed strong biological concordance with the functional roles of their corresponding domains (Supplementary Figs. C1 and C2). For example, the top-ranking genes for the shared topic 2, such as Mt3, Gpr98, Celsr1, and Nr2e1, are relevant to brain and nervous system development [20–23]. The top-ranking genes for the transcriptome private topic 10, such as Sncg, Scn9a, Nefm, and Nefl, are closely related to the trigeminal neuron clusters [24–26]. Besides, the top-ranking GASs for the epigenome private topic 3, such as Pax6, Zic2, and Hmx1, are associated with ocular development [27–29].

To validate the exclusivity of the learned private topics, we analyzed the Pearson correlation coefficients (PCCs) (Methods) between each identified topic and features from each omics dataset (Fig. 3h, Supplementary Figs. C3 and C4). In this representation, a positive log2 fold change indicates a stronger correlation with transcriptomics, while a negative value signifies a stronger correlation with epigenomics. This analysis highlights the degree to which each topic is specific to either transcriptomics or epigenomics data. The results demonstrate that all epigenome private topics exhibit a stronger correlation with epigenomics data, and all transcriptome private topics, except for the transcriptome private topic 4, show a stronger correlation with transcriptomics data.

### SpaMV facilitates identification of key developmental genes from spatial H3K27ac-H3K27me3 mouse embryo data

We applied SpaMV to another mouse embryo dataset, which involves co-profiling spatial histone modifications H3K27ac (an active marker for gene expression) and H3K27me3 (a repressive marker for gene expression) [30]. We converted the chromatin accessibility data for each histone modification into GAS (Methods), employing them as inputs for all algorithms under evaluation.

Compared with the manual annotation (Fig. 4a), SpaMV successfully identified all major anatomical structures, including the liver (cluster 16), heart (cluster 17), spine (cluster 18), and spinal cord (cluster 2) (Fig. 4b). Notably, SpaMV was the only algorithm capable of clearly separating the jaw (cluster 1), tongue (cluster 6), and nose (cluster 15) regions. Furthermore, only SpaMV and CellCharter were able to distinguish the basiocipital bone region (SpaMV cluster 19). Quantitative evaluation using the Jaccard similarity score further supports SpaMV’s performance (Fig. 4c-d). When considering both modalities, SpaMV achieved the highest Jaccard similarity (0.546), surpassing COSMOS (0.478), MISO (0.457), CellCharter (0.383), SpatialGlue (0.382), and SMOPCA (0.293). Overall, SpaMV demonstrated superior integrative performance across modalities.

**Fig. 4:**
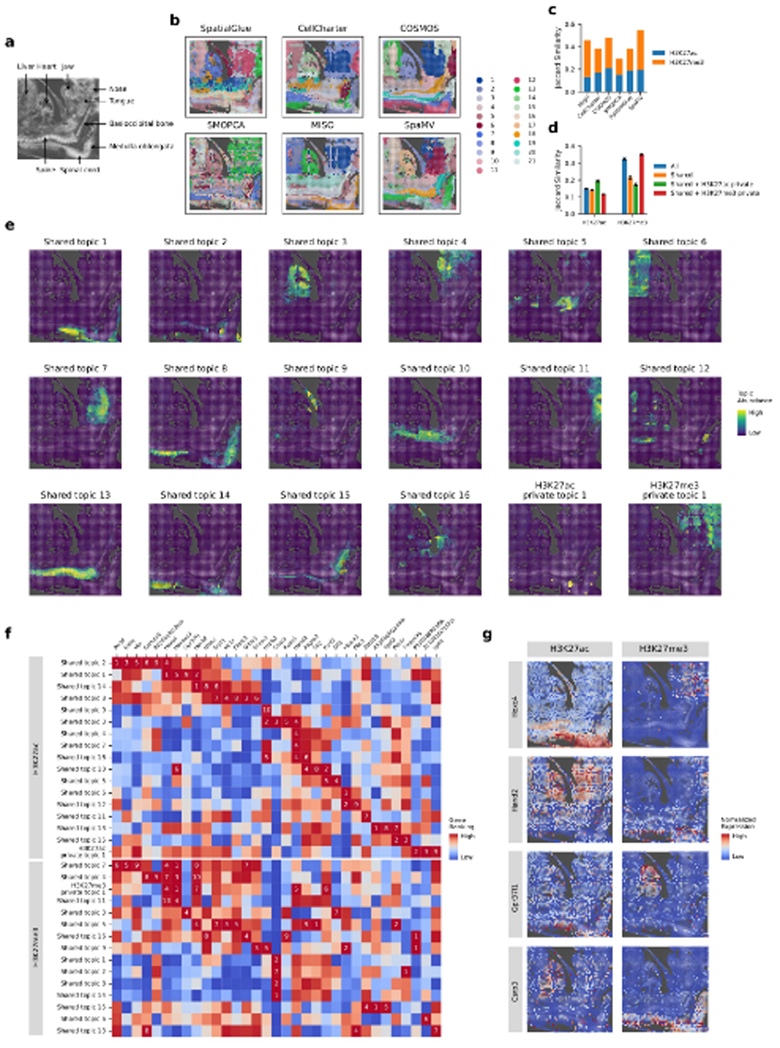
Interpretable SpaMV analysis of spatial H3K27ac-H3K27me3 mouse embryo data. **a**, Image and manual annotation of the mouse embryo data. **b**, Domain clustering results from six spatial multi-omics integration algorithms: SpatialGlue, CellCharter, COSMOS, SMOPCA, MISO, and SpaMV. **c**, Jaccard similarity scores for competing algorithms and SpaMV, where the Jaccard similarity scores for SpaMV were computed using the learned shared and omics-specific private embeddings. **d**, Jaccard similarity scores for SpaMV using different embeddings. **e**, Spatial plots of the topics learned by SpaMV through its topic modeling mode. **f**, Heat map showcasing the top-ranking genes associated with different topics across various omics layers. The color and Number indicate the ranking of a gene within a topic. **g**, Spatial plots displaying selected potential developmental marker genes with H3K27ac and H3K27me3 histone modifications.

We further applied the topic modeling mode of SpaMV on this dataset and identified 16 shared topics and one private topic for each modality (Fig. 4e, Supplementary Figs. C5 to C8), suggesting a high degree of shared information between the two histone marks. The visualized results demonstrated that the learned topics closely align with the annotated regions, such as the liver (shared topic 6), spinal cord (shared topic 14), heart (shared topic 3), jaw (shared topic 4), medulla oblongata (shared topic 8), tongue (shared topic 7), and spine (shared topic 13). Interestingly, the private topic for the H3K27ac modality is located in the spinal cord region, while the private topic for the H3K27me3 modality encompasses the jaw and nose regions (Fig. 4e).

A distinctive strength of SpaMV, when applied to this multi-modality dataset of histone modifications, is its ability to automatically identify genes that exhibit differential behavior under various histone modifications (Fig. 4f-g). Specifically, for each topic, we retained the top ten ranked genes within one modality if they also appeared among the top ten genes in another topic from the opposite modality. Using this cross-modality comparison strategy, we discovered 33 genes showing differential expression across different histone modifications. Notably, this set includes two genes (Hoxc4 and Hand2), which were also highlighted in the original study [30]. Additionally, SpaMV identified novel candidate genes not previously reported, such as Gpr37l1, implicated in nervous system development [31]; Csrp3, associated with muscle development [32]; and Hoxb8, known to play a crucial role in the development of the spinal sensory system [33].

### SpaMV robustly integrates spatial transcriptome-metabolome data in ccRCC and reveals cell-type-specific signatures

We applied SpaMV to a spatial transcriptome-metabolome dataset from clear cell renal cell carcinoma (ccRCC) (Fig. 5a) [34]. The spatial domain clustering results (Fig. 5b) revealed that SpaMV achieved more distinct and spatially coherent segmentation boundaries compared to five other algorithms. Furthermore, Jaccard similarity scores confirmed that the embeddings learned by SpaMV outperformed other competitive algorithms on both modalities (Fig. 5c-d). By leveraging the domain clustering results from SpaMV alongside a single-cell reference dataset from [34], we identified four primary cell types within the tissue section (Fig. 5e). We further applied SpaMV’s topic modeling to this dataset (Fig. 5f, Supplementary Figs. C9 and C10), and associated the learned topics with cell types based on the spatial localization of their dominant topic weights, guided by the manual annotations in Fig. 5e. This analysis revealed that each cell type was associated with at least one private topic, highlighting the importance of modeling modality-specific signals to fully capture the cellular heterogeneity present in multi-omics spatial data.

**Fig. 5:**
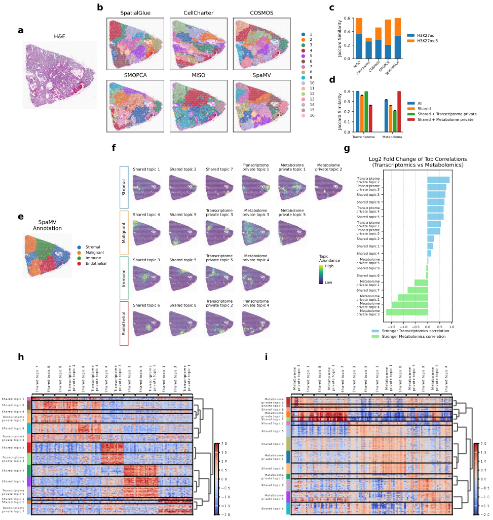
Interpretable SpaMV analysis of spatial transcriptome-metabolome clear cell renal cell carcinoma data. **a**, Hematoxylin and eosin stained image of the tissue. **b**, Domain clustering results from six spatial multi-omics integration algorithms: SpatialGlue, CellCharter, COSMOS, SMOPCA, MISO, and SpaMV. **c**, Jaccard similarity scores for competing algorithms and SpaMV, where the Jaccard similarity scores for SpaMV were computed using the learned shared and omics-specific private embeddings. **d**, Jaccard similarity scores for SpaMV using different embeddings. **e**, Manual annotation based on SpaMV domain clustering result. **f**, Spatial plots of the topics learned by SpaMV through its topic modeling mode. Each row corresponds to an annotated region. **g**, Log2 fold change of the mean Pearson correlation coefficients between transcriptome and metabolome for each topic, calculated from the top 10 correlated genes and metabolites. **h**, Heatmap of differentially expressed genes on each topic; each row represents a spatial spot, and each column corresponds to a differentially expressed gene. **i**, Heatmap of differentially expressed metabolites on each topic; each row represents a spatial spot, and each column corresponds to a differentially expressed metabolite.

Specifically, within the stromal region, the spatially variable genes associated with the transcriptome private topic 1 include key stromal cell gene markers, such as IGLC1, COL1A1, and TNC (Supplementary Fig. C9) [35]. In the immune region, the spatially variable genes associated with the transcriptome private topic 5 encompass several immunity-related gene signatures, such as GBP1 [36], TRBC2 [37], and CXCL9 [38]. We also calculated the log2 fold change of the mean top 10 PCCs for each topic across the different omics layers, as presented in Fig. 5g, Supplementary Figs. C11 and C12, which verifies that all private topics have stronger correlations with their corresponding modality.

Moreover, we further validated the assignment of learned topics by examining the differentially expressed genes and metabolites for each topic (Fig. 5h-i). To assign clusters, each spot was labeled according to the topic with the highest value in the learned topic embedding. In the transcriptomics modality, we identified four major groups that generally align well with the groupings shown in Fig. 5e. For example, the shared topics 3 and 5, along with transcriptome private topic 5, which are associated with the immune region, exhibited similar sets of differentially expressed genes. Conversely, in metabolomics, three main groups primarily cover the malignant and stromal cells. For example, the shared topic 4, shared topic 9, metabolome private topic 3, and metabolome private topic 5, which are linked to the malignant region, captured similar sets of differentially expressed metabolites.

### Accurate cell type annotation in spatial transcriptome-proteome mouse thymus data through SpaMV’s private topics

We applied SpaMV to a spatial mouse thymus data, which encompasses both transcriptomics and proteomics modalities, the latter including 19 protein markers [39]. As depicted in Fig. 6a, SpaMV effectively delineated the primary structures of the mouse thymus, including the medulla (clusters 1), the cortex (clusters 2 and 3), and the capsule (cluster 6). Quantitative evaluation using Jaccard similarity scores (Fig. 6b-c) further supports SpaMV’s performance. SpaMV achieved the highest score (0.349), substantially outperforming MISO (0.281), SpatialGlue (0.202), CellCharter (0.170), COSMOS (0.09), and spaMultiVAE (0.06).

**Fig. 6:**
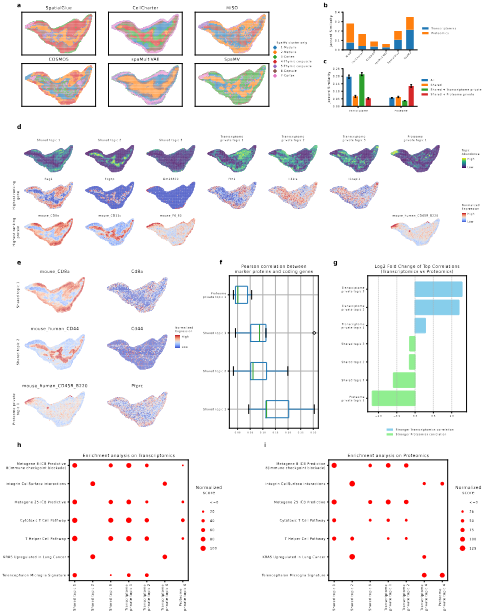
Interpretable SpaMV analysis of spatial transcriptome-proteome mouse thymus data. **a**, Domain clustering results from six spatial multi-omics integration algorithms: SpatialGlue, CellCharter, MISO, COSMOS, spaMultiVAE, and SpaMV. **b**, Jaccard similarity scores for competing algorithms and SpaMV, where the Jaccard similarity scores for SpaMV were computed using the learned shared and omics-specific private embeddings. **c**, Jaccard similarity scores for SpaMV using different embeddings. **d**, Spatial plots of the topics and associated highest ranking genes and proteins learned by SpaMV through its topic modeling mode. **e**, Spatial mappings 14 of three proteins and their corresponding coding genes. **f**, Pearson correlation coefficients between proteins and coding genes for the top five ranking proteins in shared and proteome private topics. **g**, Log2 fold change of the mean Pearson correlation coefficients between transcriptome and proteome for each topic, calculated from the top 10 correlated genes and proteins. **h**, Heatmap of pathway enrichment scores for each topic on Transcriptomics. **i**, Heatmap of pathway enrichment scores for each topic on Proteomics.

Next, we applied the interpretable topic modeling module of SpaMV to this dataset, resulting in the identification of seven distinct topics: three shared topics, three transcriptome-specific topics, and one proteome-specific topic (Fig. 6d, Supplementary Figs. C13 and C14). The shared topics aligned well with the primary structural components of the mouse thymus: the cortex (shared topic 1), medulla (shared topic 2), and capsule (shared topic 3), reflecting consistent biological signals in these regions across both modalities. Notably, shared topic 1 is characterized by high expression of the RAG1 gene, Arpp1 gene, CD8a protein, and CD4 protein, all of which are recognized as biological markers of the cortex [40–42]. Besides, shared topic 2 is distinguished by the high expression of the KRT5 gene and CD11c protein, which are markers for the medulla in the mouse thymus [43, 44]. Among the transcriptome private topics, transcriptome private topic 1, which is characterized by genes such as FTH1 and CTSL, suggests roles in active proliferation or differentiation. Transcriptome private topics 2 and 3, marked by genes like IL31RA and IL1RAPL1, are associated with thymocyte development and selection. Interestingly, despite the limited coverage of only 19 proteins, SpaMV identified a proteome-private topic. This topic features highly expressed proteins such as CD45R, IgG2a, CD19, and CD90 2, which are typical markers for B cells, suggesting possible immune cell interactions and maturation occurring in this region. Gene set variation analysis (GSVA) (Fig. 6h-i) further supported the biological relevance of the learned topics. Shared topics have similar enriched pathways on both modalities, while the proteome private topic showed strong enrichment of integrin cell surface interactions, coinciding with the function of its top-ranking genes.

Although other integration algorithms, such as SpatialGlue, MISO, and COSMOS, can also identify this proteome-specific class, their results are often difficult to interpret, which may lead to incorrect conclusions in downstream analyses. A conventional approach for annotating clusters involves identifying differentially expressed genes (DEGs) for each cluster and then comparing them to known cell type markers. As shown in Extended Data Fig. 3d, the DEGs identified for each cluster using SpaMV clustering include AY036118, Themis, and Satb1 for cluster 7. Based on these genes, one might infer that cluster 7 is enriched for T cells [45–47]. However, inspection of the spatial distribution of these DEGs (Extended Data Fig. 3f) reveals that they are not highly expressed in the cluster 7 region; instead, this region exhibits a proteomics-specific pattern (Extended Data Fig. 3a-b). In contrast, our interpretable dimension reduction approach enables users to utilize proteomics data for more accurate cell type annotation. The differentially expressed proteins for cluster 7, listed and visualized in Extended Data Fig. 3e,g, show strong spatial correspondence with cluster 7. This allows us to confidently identify the presence of B cells (CD19 [48]), plasmacytoid dendritic cells (Siglec-H [49]), and macrophages (F4/80 [50]) in this region.

The presence of this proteome private topic is likely due to the low correlation between the top-ranking proteins in this topic and their corresponding coding genes. This phenomenon is visualized in Fig. 6e, where the top-ranking proteins for shared topics 1 and 2 exhibit close spatial distribution to their coding genes, whereas the top-ranking protein for proteome-specific topic 1 displays a distinct spatial distribution from its coding gene. We further validated this by computing the PCC between proteins and coding genes for the top five ranking proteins for each shared and proteome private topic (Fig. 6f). The results indicate that the top-ranking proteins in the proteome private topic 1 generally have lower PCCs compared to those in the shared topics. Moreover, the uniqueness of the private topics was verified by examining the log2 fold change of top correlations, as shown in Fig. 6g.

## 2 Discussion

Existing spatial multi-omics integration methods often neglect modality-specific information, resulting in suboptimal spatial domain clustering and limited interpretability. To address this limitation, we developed SpaMV, a novel representation learning algorithm that explicitly disentangles shared and modality-specific information from spatial multi-omics data. Unlike previous approaches, SpaMV projects each modality into two distinct latent spaces: one capturing shared information and the other capturing modality-specific information. Leveraging these disentangled representations, SpaMV offers two types of output: (1) spatial domain clustering, which utilizes both shared and modality-specific embeddings to comprehensively characterize cellular states, and (2) interpretable dimension reduction, where structured priors promote sparsity in the latent variables, yielding interpretable topics that clarify the contribution of each modality to underlying biological processes. By adaptively utilizing these outputs, SpaMV achieves more accurate clustering and significantly enhances both the interpretability and precision of cell type annotation.

We evaluated SpaMV’s clustering and dimension reduction performance on three simulated datasets and four real-world datasets. Compared to competing algorithms such as SpatialGlue, COSMOS, CellCharter, spaMultiVAE, SMOPCA, and MISO, SpaMV achieved more accurate and interpretable results. Specifically, in the simulated datasets, some factors were designed to be modality-specific while others were designed to be common across modalities. SpaMV was the only algorithm able to successfully identify all factors, demonstrating its unique ability to preserve both shared and private information across modalities. Furthermore, SpaMV’s interpretable dimension reduction results consistently identified and clustered all factors, validating the theoretical soundness of our approach. On real-world datasets, SpaMV outperformed other integration algorithms in clustering accuracy, as measured by both supervised and unsupervised evaluation metrics. Moreover, the interpretable dimension reduction results revealed shared patterns supported by multiple modalities as well as private patterns unique to specific modalities. This information is particularly valuable for cell type annotation in multi-omics data, as it guides users to the most informative modality for annotating each cluster.

One current limitation of SpaMV is that it only considers scenarios with two modalities. When more modalities are present, information may be shared across subsets of modalities, increasing the complexity of the analysis. This challenge could be addressed by introducing additional latent variables to capture information shared among specific subsets of modalities. Although most current benchmarking efforts [51] focus on single-modality spatial transcriptomics, they highlight criteria—such as spatial alignment, biological interpretability, and geometry preservation—that any multimodal extension should retain. Similarly, recent frameworks [52], while currently limited to spatial transcriptomics, propose interpretable representation learning across multi-slice and multi-condition data, offering architectural insights relevant to multi-modal design. Additionally, some spatial multi-omics datasets include H&E-stained histology images, which can be treated as an extra modality. Despite these limitations, SpaMV remains a valuable analytical tool for spatial multi-omics integration due to its accurate and interpretable outputs. In future work, we plan to address these challenges by extending SpaMV to handle multiple modalities and incorporating histology images. We also aim to integrate single-cell multi-omics data to enable automated cell-type annotation and label transfer.

## 3 Methods

### Data preprocessing

SpaMV offers two distinct modes: a domain clustering mode and a topic modeling mode. The data preprocessing procedure differs according to the requirements of each mode.

### Domain clustering

In the domain clustering mode, SpaMV utilizes dimensionally reduced data across all input modalities to address the distributional biases present in the original data space. For spatial transcriptomics data, we initially filtered out genes expressed in fewer than 20 spots and spots with fewer than 100 expressed genes. We then selected the top 3,000 highly variable genes using the *highly variable genes* function in Scanpy [53]. Following this, normalization, log transformation, and principal component analysis (PCA) were performed using the *normalize total, log1p*, and *pca* functions in Scanpy, respectively, to prepare the final input data for SpaMV. For spatial proteomics data, we applied the centered log-ratio transformation provided by the Seurat R package [54] and used the *pca* function in Scanpy to generate the input for SpaMV. Regarding spatial epigenomics data, we encoded the data using latent semantic indexing, which we implemented independently. Finally, for spatial metabolomics data, we followed the same procedure as with spatial transcriptomics data, omitting the highly variable genes selection process.

### Topic modeling

In the topic modeling mode, SpaMV uses normalized data as input to ensure both consistency and accuracy. For any spatial omics data, we began by filtering out features expressed in less than 1% of spots and spots containing less than 1% of features as recommended by [55]. For spatial proteomics data, the centered log ratio transformation from Seurat was applied. In the case of spatial transcriptomics and spatial metabolomics data, we selected the top 1,000 highly variable features, followed by normalization and log transformation using Scanpy. For gene activity scores, which were pre-normalized, we simply filtered the top 1,000 highly variable genes using Scanpy.

### The SpaMV model architecture

The SpaMV model is an advanced variant of Variational Autoencoders (VAEs) [10], specifically designed to extract knowledge from spatial multi-omics data by disentangling shared and private information across various modalities. SpaMV provides users with two operational modes: domain clustering and topic modeling. In the domain clustering mode, SpaMV aims to preserve both the shared and private information in the learned embeddings to achieve a more accurate segmentation. In the topic modeling mode, SpaMV aims to decompose the data into shared and private topics for meaningful and interpretable dimension reduction. Both modes share a similar model architecture, but differ primarily in their decoder configurations, with the topic modeling mode incorporating an additional training phase. For a more comprehensive derivation and explanation of the SpaMV model, please refer to Spplementary Note. B.

We denote the *m*-modality spatial multi-omics data as 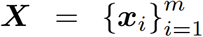. The shared latent variable, denoted as ***z***_*s*_, encapsulates information that is common across the spatial multi-omics data ***X***, while the private latent variable, ***z***_*pi*_, captures modality-specific information for the omics dataset ***x***_*i*_. We assume that the shared latent variable ***z***_*s*_ and all private latent variables 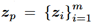 are mutually independent, therefore, the generative model for ***X*** has the form 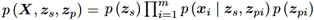. For each omics data ***x***_*i*_, we employed two distinct encoders, 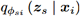 and 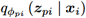, to extract ***z***_*s*_ and ***z***_*pi*_ from ***x***_*i*_, respectively. Additionally, we used a decoder 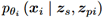 to reconstruct ***x***_*i*_ from the concatenated latent variables [***z***_*s*_, ***z***_*pi*_]. By approximating the posterior of ***z***_*s*_ by a mixture of expert operator, i.e., 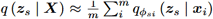, a naive evidence lower bound (ELBO) for SpaMV can be formulated as follows:

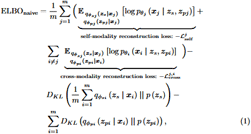

where *p* (***z***_*s*_) = *p* (***z***_*pi*_) = *N* (0, *I*). In Equation 1, the first term, referred to as the self-modality reconstruction loss 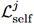, utilizes ***z***_*s*_ and ***z***_*pj*_ inferred from the same omics *j* to reconstruct ***x***_*j*_. Conversely, the second term, referred to as the cross-modality reconstruction loss 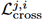, involves using ***z***_*s*_ inferred from omics *j* and ***z***_*pi*_ from omics *i* to reconstruct ***x***_*i*_. The final two terms represent the traditional KL divergence loss for the latent variables ***z***_*s*_ and ***z***_*p*_.

When this naive ELBO approaches 0, it ensures that the inferred posterior private latent variable encapsulates all modality-specific private information contained in ***x***_*i*_. This is because, during the cross-modality reconstruction, the shared latent representation input for the decoder 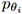 is derived from other modalities which contain no private information about ***x***_*i*_. As a result, to minimize 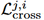, the inferred posterior 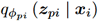 must encode all omics-*i* specific private information from ***x***_*i*_ to ***z***_*pi*_. However, this ELBO does not guarantee that: 1) the inferred shared latent variable contains all shared information, 2) no shared information leaks into the inferred private latent variable, and 3) no private information leaks into the inferred shared latent variable. To resolve these issues, we take the following three modifications, each targeting one of these specific concerns.

### Preserving all shared information in the inferred shared latent variable

To achieve this objective, we adopted the method outlined in [56]. Specifically, during cross-modality reconstruction, we replaced the private latent variable ***z***_*pi*_ with an auxiliary variable ***r***_*i*_, which follows a standard Gaussian distribution. Then, the modified cross-modality reconstruction loss 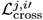 is expressed as:

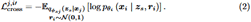

This modification ensures that all shared information is retained in the inferred shared latent variable; otherwise, the loss 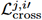 cannot be minimized. However, this approach might direct all information into the shared latent variable, leaving the private latent variables devoid of meaningful content. To address this issue, we prevented the parameters in the shared encoders from being updated during self-modality reconstruction. This adjustment reformulates the self-modality reconstruction loss as follows:

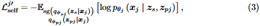

where sg (·) denotes stopping the gradient descent for the involved neural networks. This constraint ensures that the inferred private latent variable must capture all private information necessary to minimize 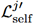.

### Preventing the leakage of shared information into inferred private latent variables

When an inferred private latent variable ***z***_*pi*_ contains shared information, it suggests non-zero mutual information between ***z***_*pi*_ and ***x***_*j*_. This mutual information can be estimated using deep neural networks, as suggested by [57]. Building on this intuition, we introduced *m* − 1 additional measurement models: 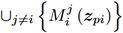 for each private latent variable ***z***_*pi*_. These models are trained to reconstruct the data ***x***_*j*_ from the private latent variable ***z***_*pi*_ by minimizing the mean squared error (MSE). The loss function for a measurement model 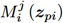 is defined as:

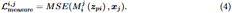

When there is no mutual information between ***z***_*pi*_ and ***x***_*j*_, the measurement model 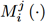 that minimizes 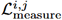 should consistently return the mean of ***x***_*j*_, irrespective of the input it receives. Thus, to ensure that no shared information is present in the inferred private latent variables, the following loss term should be minimized:

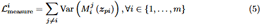

Consequently, we acquired the loss function for the SpaMV model by incorporating Equations 2, 3, and 5 with Equation 1:

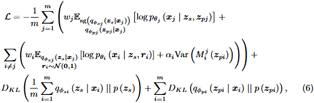

where 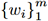 are hyperparameters that re-weight the reconstruction loss for different modalities, and 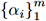 are hyperparameters that regulate the strength of the uniqueness constraint on the private latent variables. By iteratively optimizing the losses defined in Equations 4 and 6, the shared information will be effectively removed from the inferred private latent variables once both losses converge.

### Preventing the leakage of private information into inferred shared latent variables

We designed an additional training phase to remove the potential private information from the inferred shared latent variables. Since the inferred private latent variables contain all and only private information, we freeze the parameters of the private encoders and only update the parameters of the shared encoders and decoders in the second training phase. Besides, we computed the Hilbert-Schmidt independence criterion (HSIC) between the inferred shared and private latent variables to measure their dependency. HSIC is a kernel-based non-parametric statistical independence test that equals 0 if and only if the two involved sets of variables are independent [16]. Denote 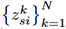 and 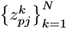 are *N* samples generated from 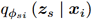 and 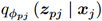, respectively. Then, the learning objective for the second training phase is:

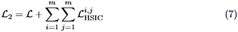

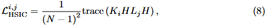

where 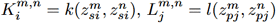 are kernel matrices with kernels *k* and *l*, respectively. 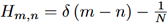 is a centering matrix. In our method, we applied Gaussian kernel to both *k* and *l*, so that 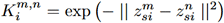, and 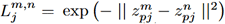.

It is worth noting that this training phase was only applied to the topic modeling mode since the presence of private information in the shared latent variable would not affect the clustering quality.

### Encoders

In the SpaMV framework, both the shared and private encoders are constructed by a graph attention network (GAT) (Spplementary Note. A) [58] which comprises *L* layers. We constructed a mixed neighboring graph, denoted as *G*_*i*_, for both the shared and private encoders of the data ***x***_*i*_. In the graph *G*_*i*_, two nodes are considered adjacent if they are either spatially neighboring or among the top *k* nearest neighbors in data space. We calculated the *k*NN graph using the *kneighbors graph* function from the scikit-learn Python package with default hyperparameters. Besides, in our experiments, we set *L* = 2 and *k* = 10 by default.

### Decoders

To effectively accomplish different tasks within the SpaMV framework, we employed two distinct types of decoders tailored to domain clustering and topic modeling to reconstruct data ***x***_*i*_ from latent variables {***z***_*s*_, ***z***_*pi*_}. In the domain clustering mode, we assumed that the data follows a Gaussian distribution, and we utilized a 2-layer multi-layer perceptron (MLP) as the decoder *D*_*i*_ for each type of spatial omics data ***x***_*i*_ to reconstruct its mean. Thus,

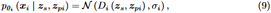

where *σ*_*i*_ is a learnable parameter that captures the standard deviation of ***x***_*i*_.

In the topic modeling mode, we adopted a parameterized model inspired by the STAMP framework [55]. Specifically, we assumed each feature *f* in the data ***x***_*i*_ follows a Gamma Poisson distribution with mean 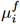 and dispersion 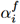 as follows:

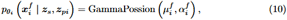

where the mean 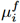 can be linearly decomposed as 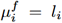· softmax (*β*_*i*_), *l*_*i*_ is the library size of ***x***_*i*_, *β*_*i*_ is the feature embedding in which each ele-ment 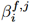 represents the contribution of feature *f* in data ***x***_*i*_ to topic ***z***_*j*_. Furthermore, to encourage the diversity of topics, STAMP applies a horseshoe prior on 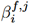, which can be defined as:

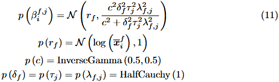

The parameters 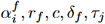 and *λ*_*f,j*_ are learned by mean-field variational inference along with the latent variables.

### Measurement models

In both domain clustering and topic modeling modes, we employed a 2-layer MLP for all measurement models. Considering the characteristics of the input data, we chose the softplus function as the final activation function layer for the domain clustering mode, and the softmax function for the topic modeling mode. Furthermore, in the topic modeling mode, we adjusted the output of the measurement model by multiplying it by the library size of the reconstructed data. This adjustment is because the inferred topics do not contain information about the absolute magnitude of the reconstructed data.

### Model training techniques

To ensure the stability of the SpaMV model, we began training the measurement models after a predetermined number of iterations. This approach ensures that the inferred private variables contain meaningful information when training the measurement models. We set the starting iteration number at 200 for the domain clustering mode and 10 for the topic modeling mode. Besides, to prevent convergence to a local optimum, we reinitialized the measurement models every 100 iterations. Each time the measurement models were initialized, we trained them for 100 iterations. The pseudocode for the SpaMV training process is outlined below:

#### Algorithm 1

The SpaMV training process

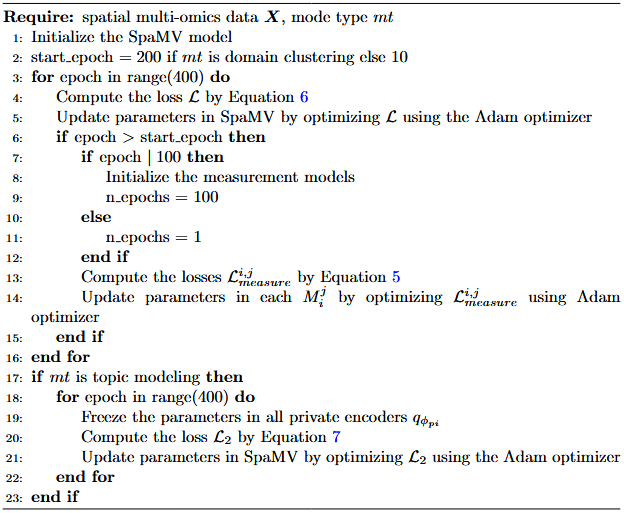

### Post-process in topic modeling

Since SpaMV is designed to simultaneously preserve both modality-specific and shared information, some background noise may occasionally be captured in the learned representations. To address this, we implemented a post-processing step in the topic modeling mode to retain only biologically meaningful dimensions within both shared and private embeddings. Specifically, we calculate Moran’s I z-score for each dimension in the learned embeddings. If the minimum z-score falls below a user-defined threshold (default: 0.3), we apply K-Means clustering to partition the dimensions into two clusters based on their z-scores. Dimensions belonging to the cluster with lower z-scores are considered noisy topics and are pruned from the results. Additionally, dimensions with low standard deviation (default: 1) are regarded as background constants without biological significance and are also removed.

### Evaluation metrics

In our domain clustering tasks, we utilized six supervised metrics: the adjusted Rand index, mutual information, adjusted mutual information, normalized mutual information, homogeneity, and the V-measure. These metrics were computed using the scikit-learn Python package. Additionally, we employed the Jaccard similarity as an unsupervised metric to assess the similarity between the learned embeddings and the original data. To compute the Jaccard similarity, we first identified two sets of *k* nearest neighbors for each spot, one based on its learned embeddings and the other on its original data. We then determined the overlapping proportion between the intersection and the union of these neighboring sets. Finally, we averaged this proportion across all spots to derive the Jaccard similarity score.

In multi-omics topic modeling tasks, we designed a new metric, called fold change of top correlations, to evaluate if a learned topic is more similar to one modality than another. Given two omics data ***x***_1_ and ***x***_2_ and a learned topic ***z***_*k*_, we first compute the Pearson correlation between the topic ***z***_*k*_ and every feature in ***x***_1_ and ***x***_2_. Then, we preserve the top *m* correlations for features in ***x***_1_, i.e., 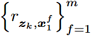, and top *n* correlations for ***x***, i.e., ***x***, i.e., 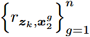, where the value of *m* and *n* depends on the modality. For Proteomics data, we preserved the top 5 Pearson correlations, and for other omics data, we preserved the top 10. Finally, the fold change of top correlation for topic ***z***_*k*_, i.e., *d* (***z***_*k*_; ***x***_1_, ***x***_2_) is defined as:

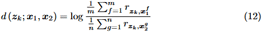

### Simulated data generation process

We followed the simulation generation process described in [59]. Specifically, we constructed a 36 *×* 36 location grid, with each representing a spot in spatial multi-omics data. These 1296 spots were divided into 9 unique 6 *×* 6 square sub-zones, each representing a distinct spatial domain. We then allocated these domains to private factors, shared factors, or a background factor as depicted in Fig. 2a, Extended Data Fig. 1a, and Extended Data Fig. 2a. Subsequently, each feature was randomly assigned to one of the shared or private factors present in its respective omics dataset. For Omics 1, we generated 1,000 features using a zero-inflated negative binomial (ZINB) distribution, while for Omics 2, we generated 100 features using a negative binomial (NB) distribution. A feature is defined as ‘active’ at a spot if it is associated with a factor assigned to that spot. For Omics 2, if a feature is active at a spot, its value is sampled with a mean of 20.2 and dispersion of 10; otherwise, the value is sampled with a mean of 0.2 and dispersion of 10. We use the same strategy to sample data in Omics 1, with an additional zero-inflation probability of 0.5. Furthermore, if a spot is designated as a background factor, there is a 5% chance of assigning a non-zero value for all features at that spot.

### Computation of gene activity score

For generating gene activity scores, we followed the tutorial from spatial-Mux-seq [30] using the createArrowFiles function from ArchR (version: 1.0.3) [60]. Seed is set to 1, and 8 threads are used with set.seed and addArchRThreads. The default genome is set to mm10 using the addArchRGenome function.

### Computation of pathway enrichment score

We computed the pathway enrichment score based on the Gene Set Variation Analysis (GSVA) package [61] with gene sets from the MSigDB database [62]. First, we assigned a topic annotation to each spot based on the result from SpaMV. Then we selected the common genes between the proteomics and transcriptomics datasets to get the subsets of the two omics layers. For each omics layer, GSVA was performed using the m2 gene sets with a Gaussian kernel and a minimum gene set size threshold of 3, outputting a matrix of enrichment scores for each pathway across all spots. We normalized the scores to the (0, 1) range for every pathway and computed the log2 fold change of each topic according to the topic annotation. Finally, we plotted dotplots for the two omics layers.

## Data availability

The mouse embryo transcriptome-epigenome multi-omics data is available at the Gene Expression Omnibus repository with the accession code GSE205055. The mouse embryo H3K27ac-H3K27me3 multi-omics data is available at the Gene Expression Omnibus repository with the accession code GSE263333. The clear cell renal cell carcinoma transcriptome-metabolome multi-omics data is available at the figshare database (https://doi.org/10.6084/m9.figshare.24599295). The mouse thymus transcriptome-proteome multi-omics data is from BGI. The simulated data and all preprocessed data are available at Zenodo (https://zenodo.org/records/16436314).

## Code availability

The source code of SpaMV is available at Github with link https://github.com/ericcombiolab/SpaMV with documentation https://spamv-tutorials.readthedocs.io/en/latest/.

## Acknowledgements

We thank all the participants for their significant contribution to this study. The design of the study and the collection, analysis and interpretation of the data were supported by the Young Collaborative Research grant (No. C2004-23Y) and HMRF grant (No. 11221026).

---

**Extended Data Figure 1:**
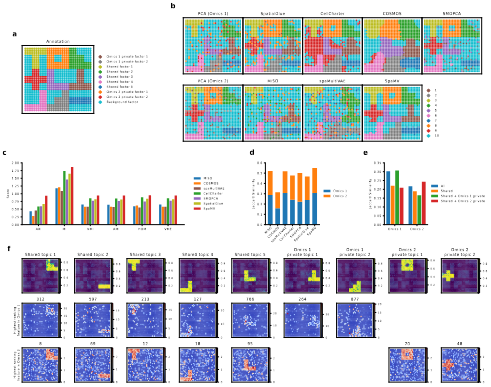
SpaMV accurately decomposes shared and private features in simulation 2. **a**, Ground truth of the simulated data 2. **b**, Spatial plots of clustering results: first column shows the mclust clustering on PCA embeddings from Omics 1 and Omics 2; subsequent columns display clustering results from SpatialGlue, CellCharter, COSMOS, SMOPCA, MISO, spaMultiVAE, and SpaMV. **c**, Quantitative evaluation of the evaluated algorithms using six supervised metrics. **d**, Jaccard similarity scores for competing algorithms and SpaMV, where SpaMV utilizes the learned shared embedding and omics-specific private embeddings. **e**, Jaccard similarity scores for SpaMV using different embeddings. **f**, Spatial plot of the learned topics and the highest ranking features identified by SpaMV.

**Extended Data Figure 2:**
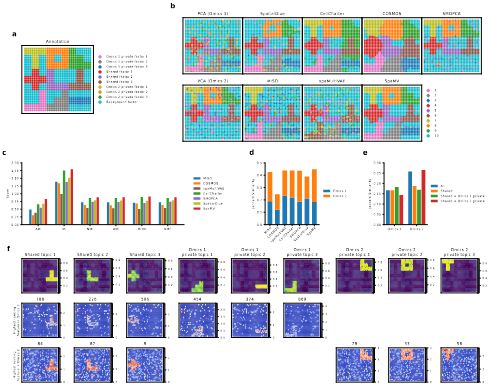
SpaMV accurately decomposes shared and private features in simulation 3. **a**, Ground truth of the simulated data 3. **b**, Spatial plots of clustering results: first column shows the mclust clustering on PCA embeddings from Omics 1 and Omics 2; subsequent columns display clustering results from SpatialGlue, CellCharter, COSMOS, SMOPCA, MISO, spaMultiVAE, and SpaMV. **c**, Quantitative evaluation of the evaluated algorithms using six supervised metrics. **d**, Jaccard similarity scores for competing algorithms and SpaMV, where SpaMV utilizes the learned shared embedding and omics-specific private embeddings. **e**, Jaccard similarity scores for SpaMV using different embeddings. **f**, Spatial plot of the learned topics and the highest ranking features identified by SpaMV.

**Extended Data Figure 3:**
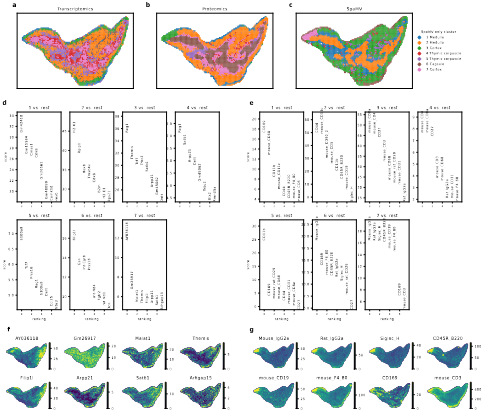
A case study for the mouse embryo data. **a**, Spatial plot of clustering result based on the Transcriptomics data, **b**, Spatial plot of clustering result based on the Proteomics data, **c**, Spatial plot of clustering result from SpaMV. **d**, Top 8 differentially expressed genes for each cluster in the SpaMV clustering result, **e**, Top 8 differentially expressed proteins for each cluster in the SpaMV clustering result, **f**, Spatial plots of the top 8 differentially expressed genes for cluster 7 in the SpaMV result, **g**, Spatial plots of the top 8 differentially expressed proteins for cluster 7 in the SpaMV result.

