## Supplementary Notes and Figures for "Interpretable spatial multi-omics data integration and dimension reduction with SpaMV"

#### Appendix A Graph attention network

Given a dataset $\boldsymbol{x} = \{\boldsymbol{x}_i\}_{i=1}^{N}$ with $N$ features, and a graph $G$ in which each node is a feature in $\boldsymbol{x}$. A graph attention network uses an attention mechanism to compute a representation of the features from the graph-structured data. Specifically, for a feature $i$ in the dataset $\boldsymbol{x}$, its node representation $h_i^l$ at layer $l$ in an $L$-layer GAT is expressed as follows:

$$h_i^l = \mathrm{ReLU}\left(\sum_{j \in \mathcal{N}_i} \alpha_{i,j} W h_i^{l-1}\right) \tag{A1}$$

$$\alpha_{i,j} = \frac{\exp\left(\mathrm{LeakyReLU}\left(a^T\left[W h_i^{l-1} \mid\mid W h_j^{l-1}\right]\right)\right)}{\sum_{k \in \mathcal{N}_i} \exp\left(\mathrm{LeakyReLU}\left(a^T\left[W h_i^{l-1} \mid\mid W h_k^{l-1}\right]\right)\right)}, \tag{A2}$$

where $h_i^0 = \boldsymbol{x}$, $a$ and $W$ are learnable parameters, $\mathcal{N}_{\rangle}$ is the neighbouring nodes of $\boldsymbol{x}_i$ in $G$, and $||$ is the concatenation operation.

#### Appendix B Derivation of SpaMV

The SpaMV model is a variational inference approach designed to disentangle private and shared information within spatial multiomics data. Suppose that the spatial multiomics data $\boldsymbol{X} = \{\boldsymbol{x}_1, \ldots, \boldsymbol{x}_m\}$ contains $m$ modalities, the main assumption of SpaMV is that each omics data set, $\boldsymbol{x}_i$, is generated by a combination of shared latent variables, $\boldsymbol{z}_s$, which capture information common to multimodal spatial omics, and private latent variables, $\boldsymbol{z}_{pi}$, which encapsulate modality-specific information that is not observable in other omics data sets. For simplicity, this paper considers two omics datasets, although the model has the potential to be extended to accommodate a higher number of modalities.

Given our primary assumption, analogous to other Variational Autoencoders (VAEs) [1], SpaMV aims to approximate the true underlying data generation process by a generative model $p_\Theta(\boldsymbol{X}, \boldsymbol{z})$ in the following form:

$$\boldsymbol{X} \sim \int p_\Theta(\boldsymbol{X}, \boldsymbol{z})\, d\boldsymbol{z}$$

$$= \int p\left(\boldsymbol{z}_s\right) \prod_{i=1}^{m} p_{\theta_i}\left(\boldsymbol{x}_i \mid \boldsymbol{z}_s, \boldsymbol{z}_{pi}\right) p\left(\boldsymbol{z}_{pi}\right) d\boldsymbol{z}, \tag{B3}$$

where $\Theta = \cup_{i=1}^{m} \{\theta_i\}$ represents the parameters in the generative neural network $p_\Theta(\boldsymbol{X}, \boldsymbol{z})$, and $\boldsymbol{z} = \cup_{i=1}^{m} \{\boldsymbol{z}_{pi}\} \cup \{\boldsymbol{z}_s\}$.

The objective of SpaMV is to maximize the likelihood of data, represented by $\log p_\theta(\boldsymbol{X})$. Due to the intractability of the posterior $p_\Theta(\boldsymbol{z} \mid \boldsymbol{X})$, we substitute it with an inference neural network $q_\Phi(\boldsymbol{z} \mid \boldsymbol{X}) = q_{\phi_{\boldsymbol{z}_s}}(\boldsymbol{z}_s \mid \boldsymbol{X}) \prod_{i=1}^{m} q_{\phi_{\boldsymbol{z}_{pi}}}(\boldsymbol{z}_{pi} \mid \boldsymbol{x}_i)$, parameterized by $\Phi = \cup_{i=1}^{m} \left\{\phi_{\boldsymbol{z}_{pi}}\right\} \cup \{\phi_{\boldsymbol{z}_s}\}$. This substitution allows us to derive the evidence lower bound (ELBO) as follows:

$$\begin{aligned}
\log p_\Theta(\boldsymbol{X}) &= \log \int p\left(\boldsymbol{z}_s\right) \prod_{i=1}^{m} p_{\theta_i}\left(\boldsymbol{x}_i \mid \boldsymbol{z}_s, \boldsymbol{z}_{pi}\right) p\left(\boldsymbol{z}_{pi}\right) d\boldsymbol{z} \\
&\geq \mathbb{E}_{\substack{q_{\phi_s}(\boldsymbol{z}_s \mid \boldsymbol{X}) \\ \prod_{i=1}^{m} q_{\phi_{pi}}(\boldsymbol{z}_{pi} \mid \boldsymbol{x}_i)}} \left[\log \frac{p\left(\boldsymbol{z}_s\right)}{q_{\phi_s}\left(\boldsymbol{z}_s \mid \boldsymbol{X}\right)} \prod_{i=1}^{m} p_{\theta_i}\left(\boldsymbol{x}_i \mid \boldsymbol{z}_s, \boldsymbol{z}_{pi}\right) \frac{p\left(\boldsymbol{z}_{pi}\right)}{q_{\phi_{pi}}\left(\boldsymbol{z}_{pi} \mid \boldsymbol{x}_i\right)}\right] \\
&= \mathbb{E}_{\substack{q_{\phi_s}(\boldsymbol{z}_s \mid \boldsymbol{X}) \\ \prod_{i=1}^{m} q_{\phi_{pi}}(\boldsymbol{z}_{pi} \mid \boldsymbol{x}_i)}} \left[\sum_{i=1}^{m} \log p_{\theta_i}\left(\boldsymbol{x}_i \mid \boldsymbol{z}_s, \boldsymbol{z}_{pi}\right)\right] - \\
&\qquad D_{KL}\left(q_{\phi_s}\left(\boldsymbol{z}_s \mid \boldsymbol{X}\right) || \, p\left(\boldsymbol{z}_s\right)\right) - \sum_{i=1}^{m} D_{KL}\left(q_{\phi_{pi}}\left(\boldsymbol{z}_{pi} \mid \boldsymbol{x}_i\right) || \, p\left(\boldsymbol{z}_{pi}\right)\right)
\end{aligned} \tag{B4}$$

To enhance the computational efficiency of our model, we further approximate the posterior of $\boldsymbol{z}_s$ using a mixture of expert (MoE) approach [2], i.e., $q_{\phi_s}(\boldsymbol{z}_s \mid \boldsymbol{X}) = \frac{1}{m} \sum_{i=1}^{m} q_{\phi_{si}}(\boldsymbol{z}_s \mid \boldsymbol{x}_i)$. Consequently, the ELBO can be expressed as follows:

$$\begin{aligned}
\log p_\Theta(\boldsymbol{X}) \geq \frac{1}{m} \sum_{j=1}^{m} \Bigg( & \underbrace{\mathbb{E}_{\substack{q_{\phi_{sj}}(\boldsymbol{z}_s \mid \boldsymbol{x}_j) \\ q_{\phi_{pj}}(\boldsymbol{z}_{pj} \mid \boldsymbol{x}_j)}} \left[\log p_{\theta_j}\left(\boldsymbol{x}_j \mid \boldsymbol{z}_s, \boldsymbol{z}_{pj}\right)\right]}_{\text{self-modality reconstruction: } \mathcal{L}_{\text{self}}^{j}} + \\
& \sum_{i \neq j} \underbrace{\mathbb{E}_{\substack{q_{\phi_{sj}}(\boldsymbol{z}_s \mid \boldsymbol{x}_j) \\ q_{\phi_{pi}}(\boldsymbol{z}_{pi} \mid \boldsymbol{x}_i)}} \left[\log p_{\theta_i}\left(\boldsymbol{x}_i \mid \boldsymbol{z}_s, \boldsymbol{z}_{pi}\right)\right]}_{\text{cross-modality reconstruction: } \mathcal{L}_{\text{cross}}^{j,i}} \Bigg) - \\
& D_{KL}\left(\frac{1}{m} \sum_{i=1}^{m} q_{\phi_{si}}\left(\boldsymbol{z}_s \mid \boldsymbol{x}_i\right) || \, p\left(\boldsymbol{z}_s\right)\right) - \\
& \sum_{i=1}^{m} D_{KL}\left(q_{\phi_{pi}}\left(\boldsymbol{z}_{pi} \mid \boldsymbol{x}_i\right) || \, p\left(\boldsymbol{z}_{pi}\right)\right)
\end{aligned} \tag{B5}$$

In Equation B5, we define the self-modality reconstruction as the process of reconstructing data $\boldsymbol{x}_j$ using the shared latent variables $\boldsymbol{z}_s$ derived from the reconstructed data itself, denoted by $\boldsymbol{z}_s \sim q_{\phi_{si}}(\boldsymbol{z}_s \mid \boldsymbol{x}_j)$. Conversely, cross-modality reconstruction involves reconstructing data $\boldsymbol{x}_i$ using shared latent variables $\boldsymbol{z}_s$ extracted from different data, represented as $\boldsymbol{z}_s \sim q_{\phi_{sj}}\left(\boldsymbol{z}_s \mid \boldsymbol{x}_j\right)$. When this naive ELBO approaches 0, it ensures that the inferred posterior private latent variable encapsulates all modality-specific private information contained in $\boldsymbol{x}_i$. This is because, during the cross-modality reconstruction, the shared latent representation input for the decoder $p_{\theta_i}$ is derived from other modalities which contain no private information about $\boldsymbol{x}_i$. As a result, to minimize $\mathcal{L}_{\text{cross}}^{j,i}$, the inferred posterior $q_{\phi_{pi}}\left(\boldsymbol{z}_{pi} \mid \boldsymbol{x}_i\right)$ must encode all omics-$i$ specific private information from $\boldsymbol{x}_i$ to $\boldsymbol{z}_{pi}$. However, this ELBO does not guarantee that: 1) the inferred shared latent variable contains all shared information, 2) no shared information leaks into the inferred private latent variable, and 3) no private information leaks into the inferred shared latent variable. To resolve these issues, we take the following three modifications, each targeting one of these specific concerns.

1. Preserving shared information in the inferred shared latent variable

Since both reconstruction processes rely on private latent variables encoded from target data, there is a risk that all meaningful information is captured solely in the private latent space, rendering the shared latent variables insignificant [3]. This outcome is undesirable, as it undermines the purpose of utilizing shared latent variables to capture commonalities across different data modalities. To mitigate this issue, we adopt the modification suggested in [3], wherein the private latent variables used in cross-modality reconstruction are replaced by random Gaussian variables of the same shape. As a result, the shared latent variables are compelled to capture shared information across modalities to effectively optimize the cross-modality reconstruction loss. Based on this idea, we can derive the following ELBO:

$$
\begin{aligned}
\log p_{\Theta}(\boldsymbol{X}) &= \log \int p\left(\boldsymbol{z}_s\right) \prod_{i=1}^{m} p_{\theta_i}\left(\boldsymbol{x}_i \mid \boldsymbol{z}_s\right) d\boldsymbol{z}_s \\
&\geq \mathbb{E}_{q_{\phi_s}\left(\boldsymbol{z}_s \mid \boldsymbol{X}\right)}\left[\log \frac{p\left(\boldsymbol{z}_s\right)}{q_{\phi_s}\left(\boldsymbol{z}_s \mid \boldsymbol{X}\right)} \prod_{i=1}^{m} p_{\theta_i}\left(\boldsymbol{x}_i \mid \boldsymbol{z}_s\right)\right] \\
&= \frac{1}{m} \sum_{j=1}^{m}\left(\mathbb{E}_{q_{\phi_{sj}}\left(\boldsymbol{z}_s \mid x_j\right)}\left[\log p_{\theta_j}\left(\boldsymbol{x}_j \mid \boldsymbol{z}_s\right)\right]+\sum_{i \neq j} \mathbb{E}_{q_{\phi_{sj}}\left(\boldsymbol{z}_s \mid \boldsymbol{x}_i\right)}\left[\log p_{\theta_i}\left(\boldsymbol{x}_i \mid \boldsymbol{z}_s\right)\right]\right)- \\
&\quad D_{KL}\left(q_{\phi_s}\left(\boldsymbol{z}_s \mid \boldsymbol{X}\right) \| \, p\left(\boldsymbol{z}_s\right)\right) \\
&\geq \frac{1}{m} \sum_{j=1}^{m}\left(\mathbb{E}_{\substack{q_{\phi_{sj}}\left(\boldsymbol{z}_s \mid \boldsymbol{x}_j\right) \\ q_{\phi_{pj}}\left(\boldsymbol{z}_{pj} \mid \boldsymbol{x}_j\right)}}\left[\log p_{\theta_j}\left(\boldsymbol{x}_j \mid \boldsymbol{z}_s, \boldsymbol{z}_{pj}\right) \frac{p\left(\boldsymbol{z}_{pj}\right)}{q_{\phi_{pj}}\left(\boldsymbol{z}_{pj} \mid \boldsymbol{x}_j\right)}\right]+\right. \\
&\quad \left.\sum_{i \neq j} \mathbb{E}_{q_{\phi_{sj}}\left(\boldsymbol{z}_s \mid \boldsymbol{x}_i\right)}\left[\log p_{\theta_i}\left(\boldsymbol{x}_i \mid \boldsymbol{z}_s\right)\right]\right)-D_{KL}\left(q_{\phi_s}\left(\boldsymbol{z}_s \mid \boldsymbol{X}\right) \| \, p\left(\boldsymbol{z}_s\right)\right) \\
&\geq \frac{1}{m} \sum_{j=1}^{m}\left(\mathbb{E}_{\substack{q_{\phi_{sj}}\left(\boldsymbol{z}_s \mid \boldsymbol{x}_j\right) \\ q_{\phi_{pj}}\left(\boldsymbol{z}_{pj} \mid \boldsymbol{x}_j\right)}}\left[\log p_{\theta_j}\left(\boldsymbol{x}_j \mid \boldsymbol{z}_s, \boldsymbol{z}_{pj}\right)\right]+\right.
\end{aligned}
$$

$$\sum_{\substack{i\neq j}} \mathbb{E}_{\substack{q_{\phi_{sj}}(\boldsymbol{z}_s|\boldsymbol{x}_i)\\ \boldsymbol{w}_i\sim\mathcal{N}(\boldsymbol{0},\boldsymbol{1})}} \left[\log p_{\theta_i}\left(\boldsymbol{x}_i \mid \boldsymbol{z}_s, \boldsymbol{w}_i\right)\right]\Bigg) -$$

$$D_{KL}\left(q_{\phi_s}\left(\boldsymbol{z}_s \mid \boldsymbol{X}\right) || \, p\left(\boldsymbol{z}_s\right)\right) - \sum_{i=1}^{m} D_{KL}\left(q_{\phi_{pi}}\left(\boldsymbol{z}_{pi} \mid \boldsymbol{x}_i\right) || \, p\left(\boldsymbol{z}_{pi}\right)\right) \tag{B6}$$

Equation B6 may cause the inferred private latent variables to lack essential private information since the shared variable is used in both self-modality and cross-modality reconstruction processes. To prevent this, we halted the backpropagation of parameters in the shared encoder during self-modality reconstruction, ensuring that these parameters are updated solely by the cross-modality reconstruction loss. Thus, we have the following ELBO:

$$\log p_{\Theta}\left(\boldsymbol{X}\right) \geq \frac{1}{m}\sum_{j=1}^{m}\Bigg(\mathbb{E}_{\substack{\mathrm{sg}\left(q_{\phi_{sj}}(\boldsymbol{z}_s|\boldsymbol{x}_j)\right)\\ q_{\phi_{pj}}(\boldsymbol{z}_{pj}|\boldsymbol{x}_j)}}\left[\log p_{\theta_j}\left(\boldsymbol{x}_j \mid \boldsymbol{z}_s, \boldsymbol{z}_{pj}\right)\right] +$$

$$\sum_{\substack{i\neq j}} \mathbb{E}_{\substack{q_{\phi_{sj}}(\boldsymbol{z}_s|\boldsymbol{x}_i)\\ \boldsymbol{w}_i\sim\mathcal{N}(\boldsymbol{0},\boldsymbol{1})}} \left[\log p_{\theta_i}\left(\boldsymbol{x}_i \mid \boldsymbol{z}_s, \boldsymbol{w}_i\right)\right]\Bigg) -$$

$$D_{KL}\left(q_{\phi_s}\left(\boldsymbol{z}_s \mid \boldsymbol{X}\right) || \, p\left(\boldsymbol{z}_s\right)\right) - \sum_{i=1}^{m} D_{KL}\left(q_{\phi_{pi}}\left(\boldsymbol{z}_{pi} \mid \boldsymbol{x}_i\right) || \, p\left(\boldsymbol{z}_{pi}\right)\right), \tag{B7}$$

where the operator $\mathrm{sg}\left(\cdot\right)$ means stopping gradient descent of the parameters in the involved neural networks.

2. Preventing the leakage of shared information in the inferred private latent variables

As noted in [4], when two sets of variables are independent, training a deep neural network to use one set as input to predict the other will consistently yield the mean of the predicted variables if the model is optimized using mean squared error. Inspired by this basic idea, we introduced $m-1$ measurement models, denoted as $\left\{M_i^j\right\}_{j\neq i}$ for each private variable $\boldsymbol{z}_{pi}$ to ensure their independence from data $\cup_{j\neq i}\boldsymbol{x}_j$. Specifically, each measurement model $M_i^j$ takes $\boldsymbol{z}_{pi}$ as input and aims to predict the data $\boldsymbol{x}_j$ by minimizing the mean squared error between the prediction and the observation. If the inferred private latent variable $\boldsymbol{z}_{pi}$ contains only the private information from $\boldsymbol{x}_i$, and the measurement model $M_i^j$ reaches its global optimum, then the measurement model $M_i^j$ will consistently return the mean $\overline{\boldsymbol{x}}_j$, i.e., $\mathrm{Var}\left(M_i^j\left(\boldsymbol{z}_{pi}\right)\right) \rightarrow 0$. Thus, by iteratively optimizing our main model alongside these auxiliary measurement models, we can effectively prevent the leakage of shared information into the inferred private latent variables. The loss for the measurement model $M_i^j$ is:

$$\mathcal{L}_{measure}^{i,j} = \mathrm{MSE}\left(M_i^j\left(\boldsymbol{z}_{pi}\right), \boldsymbol{x}_j\right) \tag{B8}$$

The updated loss function of the main model can be written as:

$$\mathcal{L} = -\frac{1}{m}\sum_{j=1}^{m}\Bigg(\mathbb{E}_{\substack{\mathrm{sg}\left(q_{\phi_{sj}}(\boldsymbol{z}_s|\boldsymbol{x}_j)\right)\\ q_{\phi_{pj}}(\boldsymbol{z}_{pj}|\boldsymbol{x}_j)}}\left[\log p_{\theta_j}\left(\boldsymbol{x}_j \mid \boldsymbol{z}_s, \boldsymbol{z}_{pj}\right)\right] +$$

$$\sum_{i\neq j}\mathbb{E}_{\substack{q_{\phi_{sj}}(\boldsymbol{z}_s|\boldsymbol{x}_i)\\ \boldsymbol{w}_i\sim\mathcal{N}(\mathbf{0},\mathbf{1})}}\left[\log p_{\theta_i}\left(\boldsymbol{x}_i \mid \boldsymbol{z}_s, \boldsymbol{w}_i\right)\right] + \mathrm{Var}\left(M_i^j\left(\boldsymbol{z}_{pj}\right)\right)\Bigg) +$$

$$D_{KL}\left(q_{\phi_s}\left(\boldsymbol{z}_s \mid \boldsymbol{X}\right) || \, p\left(\boldsymbol{z}_s\right)\right) + \sum_{i=1}^{m} D_{KL}\left(q_{\phi_{pi}}\left(\boldsymbol{z}_{pi} \mid \boldsymbol{x}_i\right) || \, p\left(\boldsymbol{z}_{pi}\right)\right) \tag{B9}$$

3. Preventing the leakage of private information in the inferred shared latent variables

To eliminate any potential private information from the inferred shared latent variable, we designed an additional training phase after the above processes have converged. After the main model and the measurement models have minimized Equations B9 and B8, we can guarantee that the inferred private latent variables contain all and only private information. Therefore, during this subsequent training phase, the private encoders are kept fixed, while only the parameters of the shared encoders and decoders are actively updated to promote independence between the shared and private latent variables. This selective training approach is instrumental in refining the shared latent variables, ensuring that any residual private information initially captured is effectively removed. By focusing the optimization process solely on the shared components, we facilitate a clearer distinction between shared and private information, enhancing the model's ability to represent common features across different data sources while maintaining the integrity of the private information.

To quantitatively assess and enforce the independence between the inferred shared and private latent variables, we employed the Hilbert-Schmidt Independence Criterion (HSIC) [5]. HSIC is a powerful, kernel-based non-parametric statistical test that measures the degree of dependence between two sets of variables. It evaluates to zero if and only if the two sets of variables are independent.

Let's denote $\left\{\boldsymbol{z}_{si}^{k}\right\}_{k=1}^{N}$ and $\left\{\boldsymbol{z}_{pj}^{k}\right\}_{k=1}^{N}$ as N samples drawn from the distributions $q_{\phi_{si}}\left(\boldsymbol{z}_s \mid \boldsymbol{x}_i\right)$ and $q_{\phi_{pj}}\left(\boldsymbol{z}_{pj} \mid \boldsymbol{x}_j\right)$, respectively. The learning objective for the second training phase incorporates both the original loss function $\mathcal{L}$ and a penalty term based on HSIC to minimize dependency between shared and private variables:

$$\mathcal{L}_2 = \mathcal{L} + \sum_{i=1}^{m}\sum_{j=1}^{m}\mathrm{HSIC}\left(\left\{\boldsymbol{z}_{si}^{k}, \boldsymbol{z}_{pj}^{k}\right\}_{k=1}^{N}\right). \tag{B10}$$

The HSIC between these sets of variables is computed as:

$$\mathrm{HSIC}\left(\left\{z_{si}^{k}, z_{pj}^{k}\right\}_{k=1}^{N}\right) = \frac{1}{(N-1)^2}\mathrm{trace}\left(K_i H L_j H\right), \tag{B11}$$

where $K_i^{m,n} = k(z_{si}^m, z_{si}^n)$, $L_j^{m,n} = l(z_{pj}^m, z_{pj}^n)$ are kernel matrices constructed using kernel functions $k$ and $l$, respectively. $H_{m,n} = \delta(m-n) - \frac{1}{N}$ is a centering matrix used to ensure that the kernel matrices are centered in feature space.

For computational efficiency and effectiveness, we applied the Gaussian kernel to both $k$ and $l$. This choice is motivated by the Gaussian kernel's ability to capture complex relationships between variables. Consequently, we have:

$$K_i^{m,n} = \exp\left(- \| z_{si}^m - z_{si}^n \|^2\right) \tag{B12}$$

$$L_j^{m,n} = \exp\left(- \| z_{pj}^m - z_{pj}^n \|^2\right) \tag{B13}$$

Through this carefully designed second training phase, we iteratively refine the shared latent variables, ensuring they remain free of private information.

### Appendix C Supplementary Figures

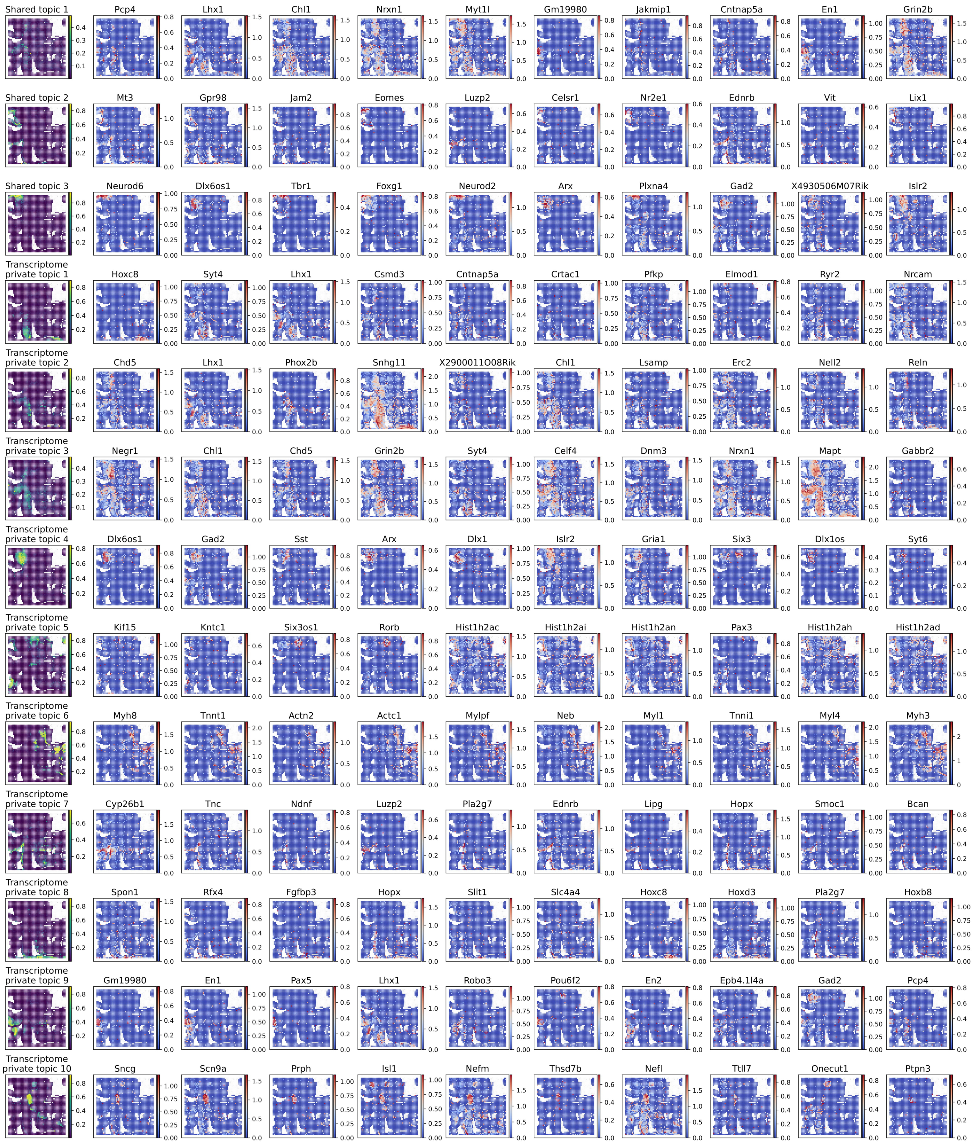


**Supplementary Figure C1**: The shared and transcriptome private topics (the first column) and their corresponding top 10 ranking genes (the second to eleventh columns) according to the learned feature module embedding for the spatial transcriptome-epigenome mouse embryo dataset.

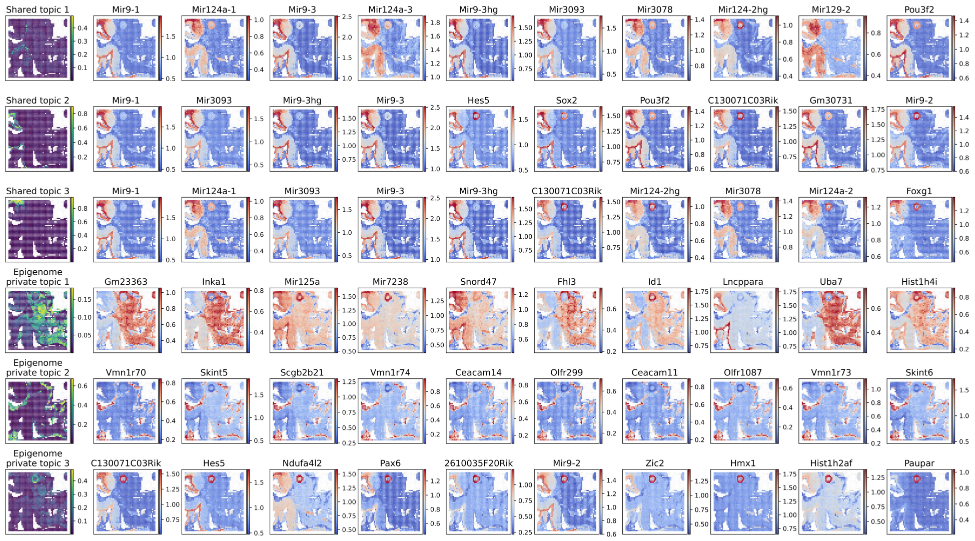


**Supplementary Figure C2**: The shared and epigenome private topics (the first column) and their corresponding top 10 ranking genes (the second to eleventh columns) according to the learned feature module embedding for the spatial transcriptome-epigenome mouse embryo dataset.

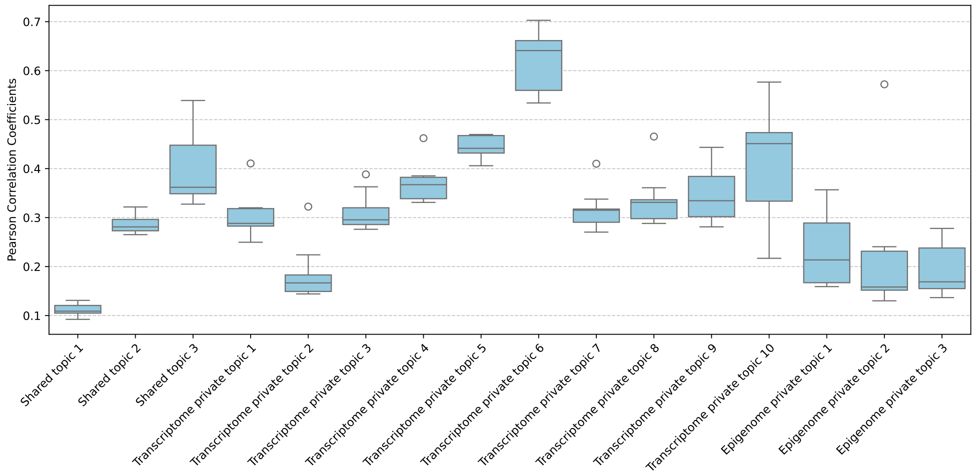


**Supplementary Figure C3**: Box plots illustrating the top 10 Pearson correlation coefficients calculated between each topic and gene from the spatial transcriptome-epigenome mouse embryo dataset.

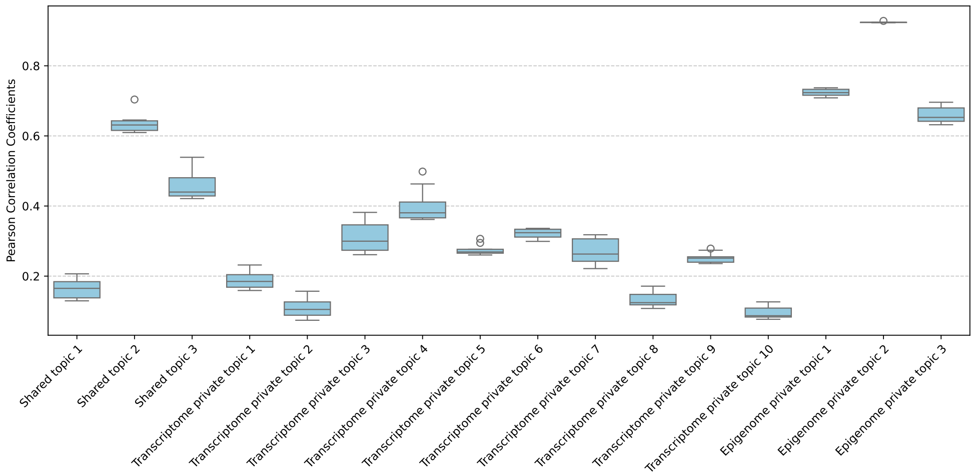


**Supplementary Figure C4**: Box plots illustrating the top 10 Pearson correlation coefficients calculated between each topic and gene derived from the gene activity score converted from the epigenomics modality in the spatial transcriptome-epigenome mouse embryo dataset.

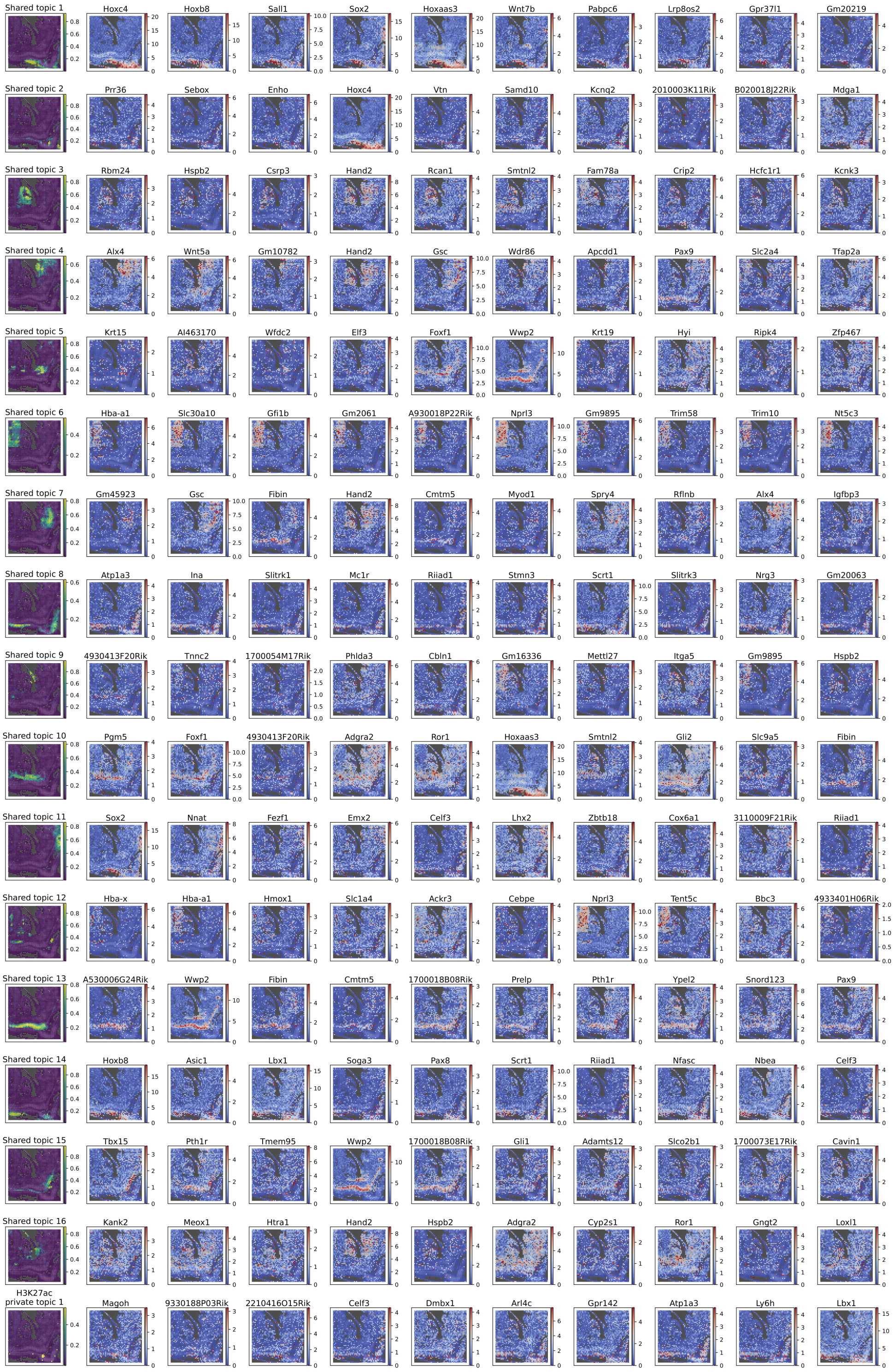


**Supplementary Figure C5**: The shared and H3K27ac private topics (the first column) and their corresponding top 10 ranking genes (the second to eleventh columns) according to the learned feature module embedding for the spatial H3K27ac-H3K27me3 mouse embryo dataset.

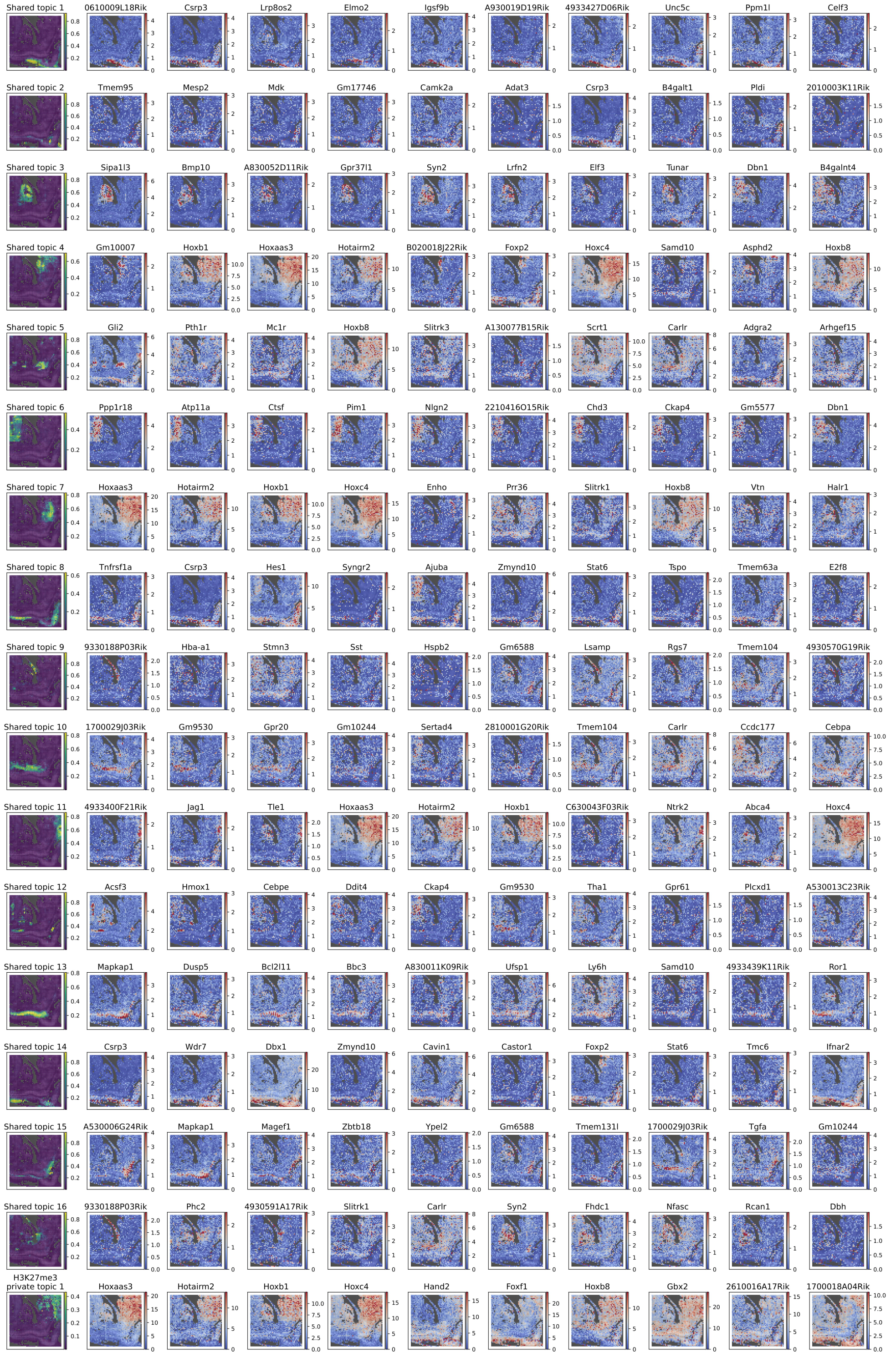


**Supplementary Figure C6**: The shared and H3K27me3 private topics (the first column) and their corresponding top 10 ranking genes (the second to eleventh columns) according to the learned feature module embedding for the spatial H3K27ac-H3K27me3 mouse embryo dataset.

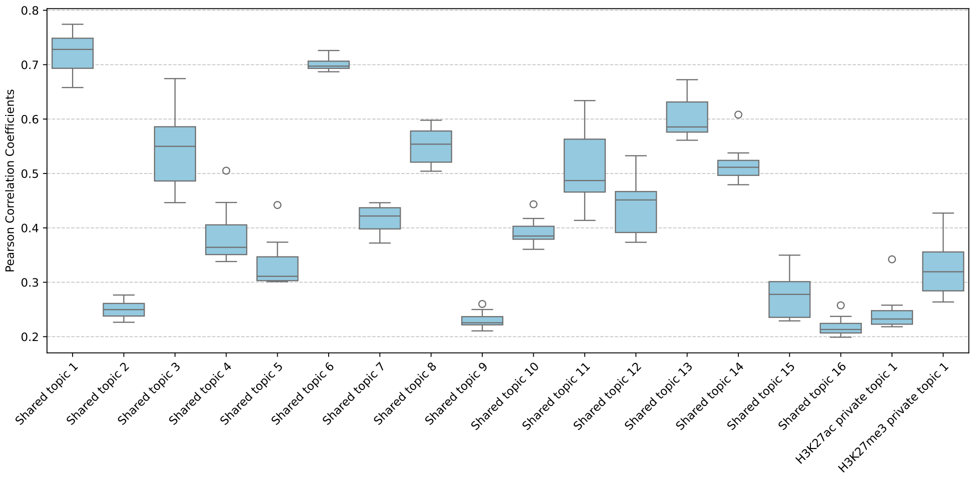


**Supplementary Figure C7**: Box plots illustrate the top 10 Pearson correlation coefficients calculated between each topic and gene derived from the gene activity score converted from H3K27ac in the spatial H3K27ac-H3K27me3 mouse embryo dataset.

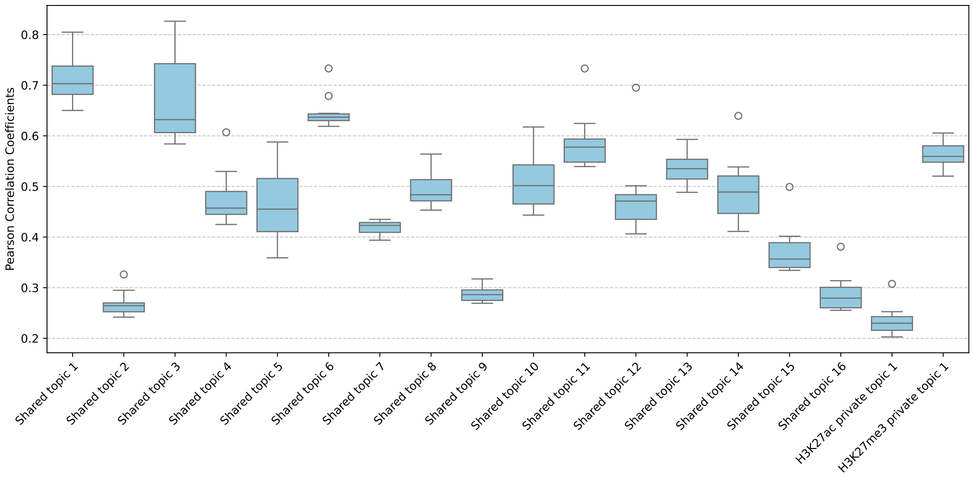


**Supplementary Figure C8**: Box plots illustrate the top 10 Pearson correlation coefficients calculated between each topic and gene derived from the gene activity score converted from H3K27me3 in the spatial H3K27ac-H3K27me3 mouse embryo dataset.

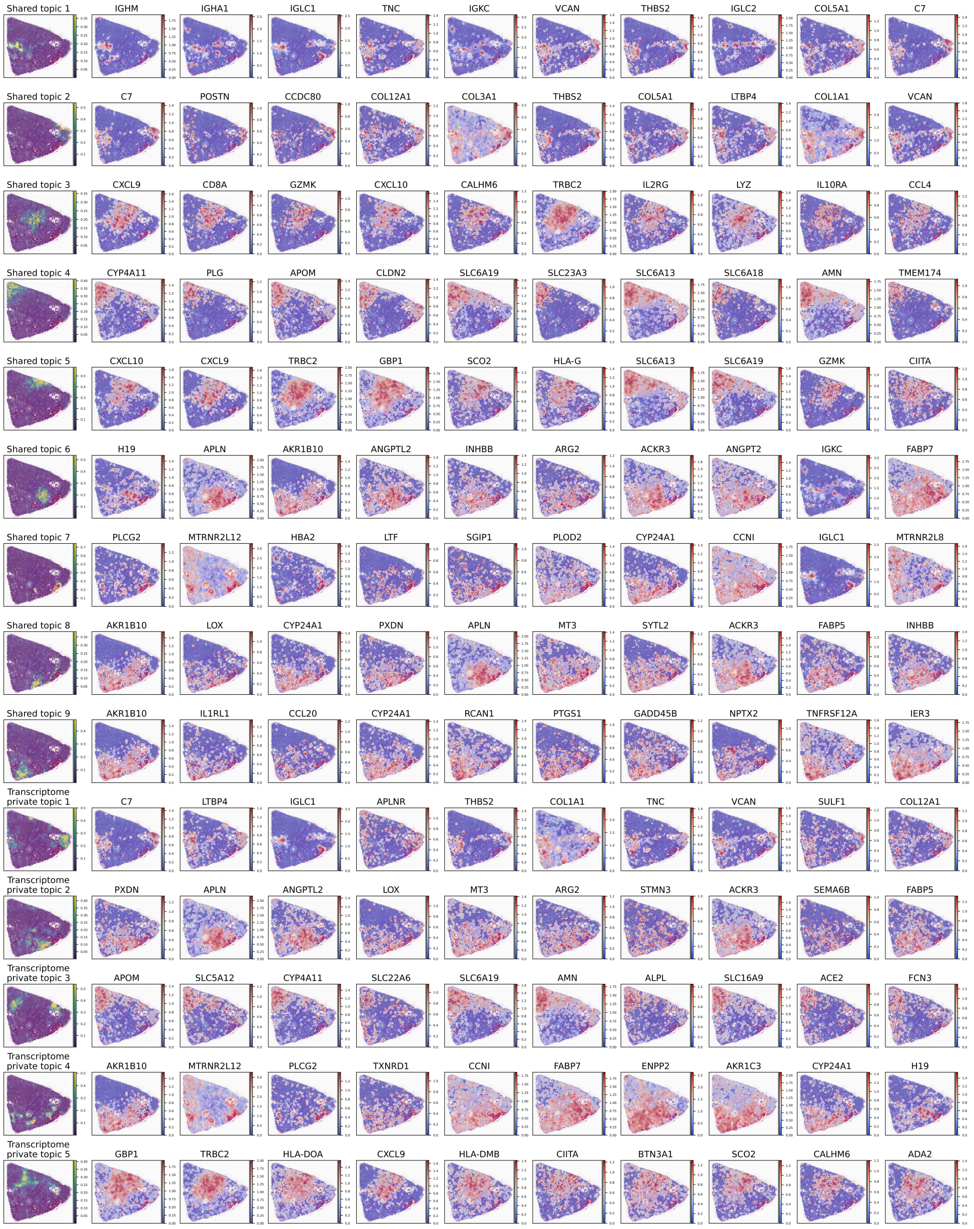


**Supplementary Figure C9**: The shared and transcriptome private topics (the first column) and their corresponding top 10 ranking genes (the second to eleventh columns) according to the learned feature module embedding for the spatial transcriptome-metabolome clear cell renal cell carcinoma dataset.

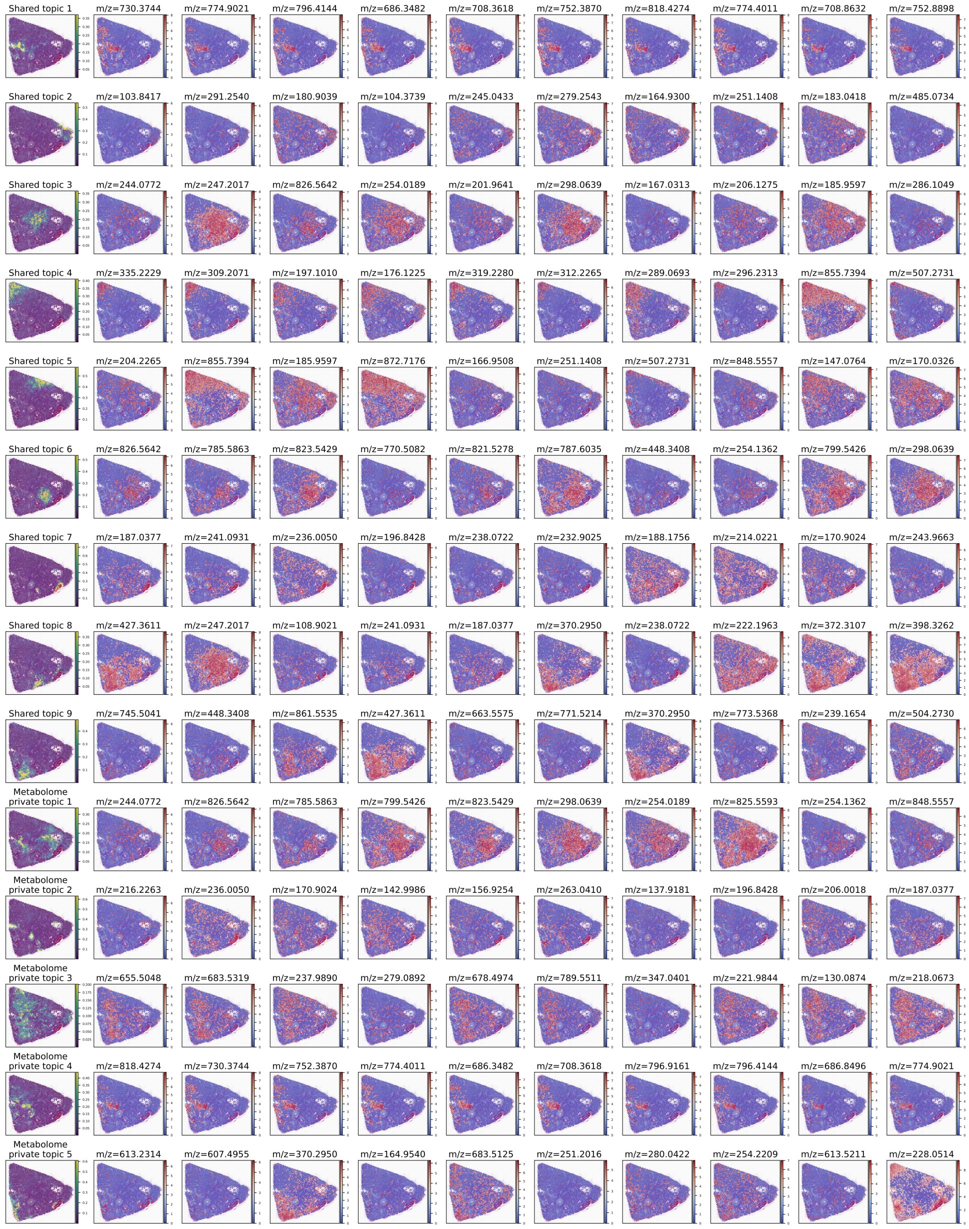


**Supplementary Figure C10**: The shared and metabolome private topics (the first column) and their corresponding top 10 ranking metabolites (the second to eleventh columns) according to the learned feature module embedding for the spatial transcriptome-metabolome clear cell renal cell carcinoma dataset.

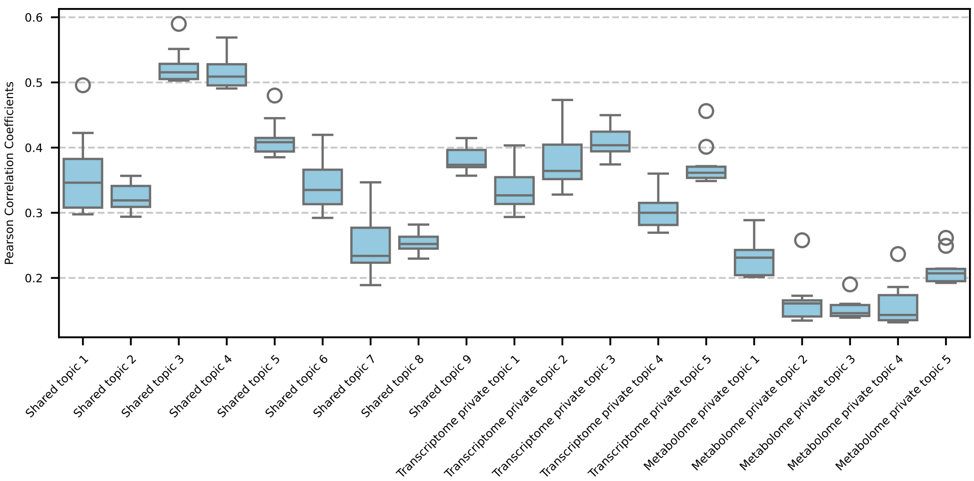


**Supplementary Figure C11**: Box plots illustrate the top 10 Pearson correlation coefficients calculated between each topic and gene in the spatial transcriptome-metabolome clear cell renal cell carcinoma dataset.

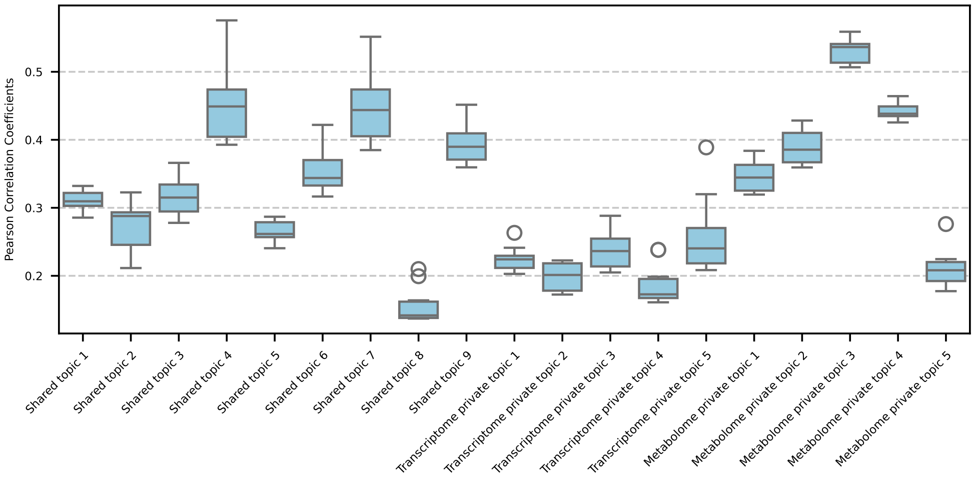


**Supplementary Figure C12**: Box plots illustrate the top 10 Pearson correlation coefficients calculated between each topic and metabolite in the spatial transcriptome-metabolome clear cell renal cell carcinoma dataset.

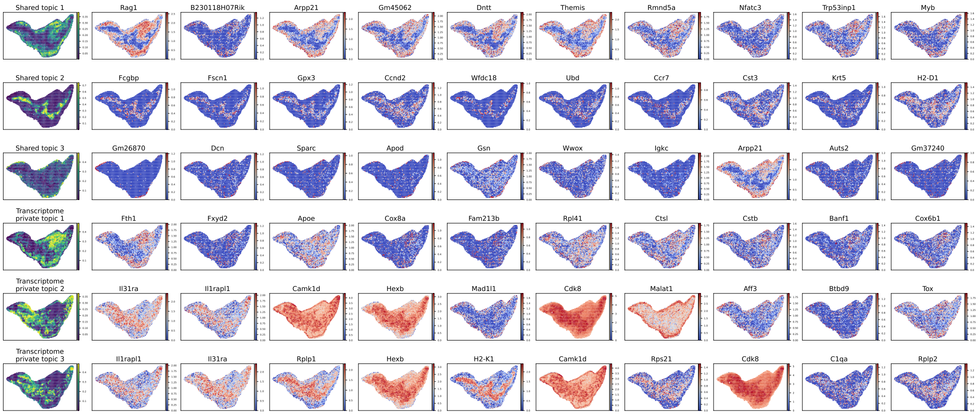


**Supplementary Figure C13**: The shared and transcriptome private topics (the first column) and their corresponding top 10 ranking genes (the second to eleventh columns) according to the learned feature module embedding for the spatial transcriptome-proteome mouse thymus dataset.

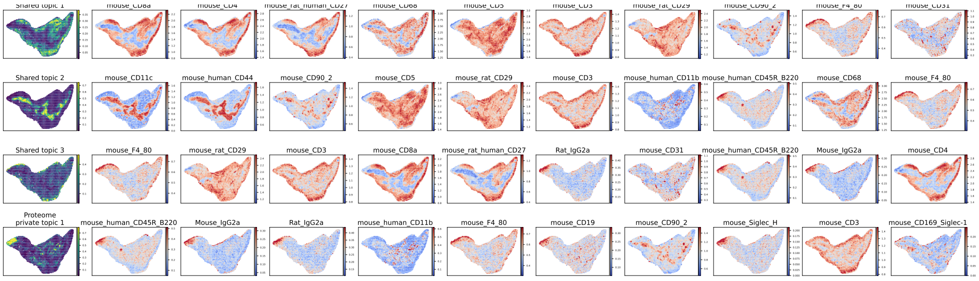


**Supplementary Figure C14**: The shared and proteome private topics (the first column) and their corresponding top 10 ranking proteins (the second to eleventh columns) according to the learned feature module embedding for the spatial transcriptome-proteome mouse thymus dataset.

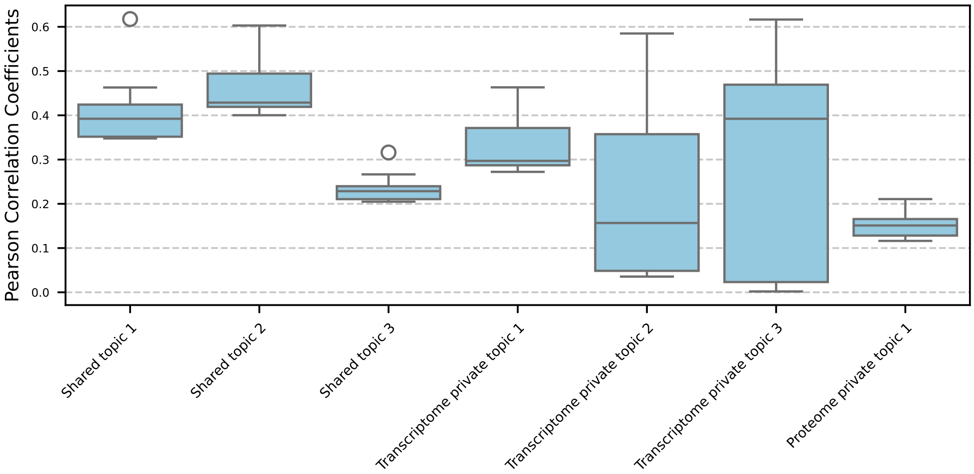


**Supplementary Figure C15**: Box plots illustrate the top 10 Pearson correlation coefficients calculated between each topic and gene in the spatial transcriptome-proteome mouse thymus dataset.

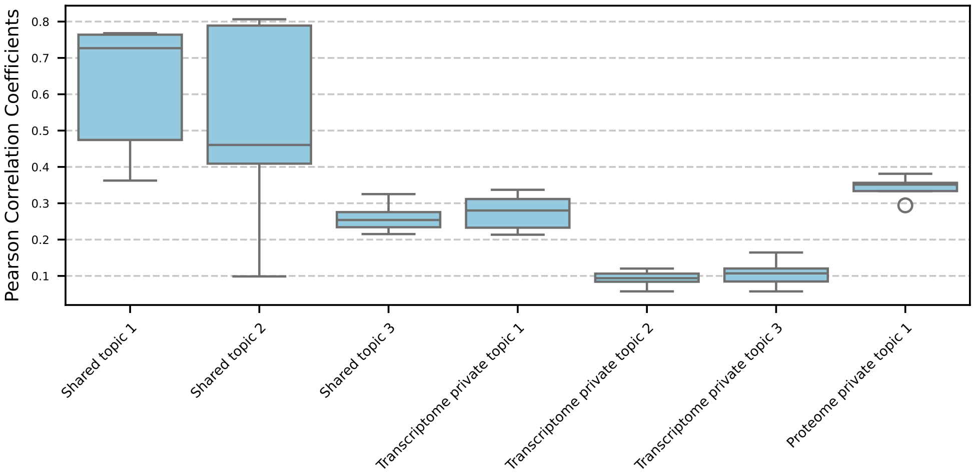


**Supplementary Figure C16**: Box plots illustrate the top 10 Pearson correlation coefficients calculated between each topic and protein in the spatial transcriptome-proteome mouse thymus dataset.
